# Prolonged partner separation erodes nucleus accumbens transcriptional signatures of pair bonding in male prairie voles

**DOI:** 10.1101/2021.07.14.452355

**Authors:** Julie M. Sadino, Xander G. Bradeen, Conor J. Kelly, Liza E. Brusman, Deena M. Walker, Zoe R. Donaldson

## Abstract

The loss of a spouse is often cited as the most traumatic event in a person’s life. However, for most people, the severity of grief and its maladaptive effects subside over time via an understudied adaptive process. Like humans, socially monogamous prairie voles (*Microtus ochrogaster*) form opposite-sex pair bonds, and upon partner separation, show stress phenotypes that diminish over time. We test the hypothesis that extended partner separation diminishes pair bond-associated behaviors and causes pair bond transcriptional signatures to erode. Pairs were cohoused for 2 weeks and then either remained paired or were separated for 48hrs or 4wks before collecting fresh nucleus accumbens tissue for RNAseq. In a separate cohort, we assessed partner preference and selective aggression at these time points, finding that these behaviors persist despite prolonged separation in both same-sex and opposite-sex paired voles. Opposite-sex pair bonding led to changes in accumbal transcription that were stably maintained while animals remained paired but eroded following prolonged partner separation. Eroded genes are associated with gliogenesis and myelination, suggesting a previously undescribed role for glia in pair bonding and loss. Further, we pioneered neuron-specific translating ribosomal affinity purification in voles. Neuronally-enriched transcriptional changes revealed dopaminergic-, mitochondrial-, and steroid hormone signaling-associated gene clusters sensitive to acute pair bond disruption and loss adaptation. Our results suggest that partner separation erodes transcriptomic signatures of pair bonding despite core behavioral features of the bond remaining intact, revealing potential molecular processes priming a vole to be able to form a new bond.

## Introduction

The death of a romantic partner is cited as one of the most traumatic experiences in a person’s life and results in grief and corresponding indicators of mental and physiological distress (*1, 2*). However, for the majority of people acute grief subsides within approximately six months as the bereaved integrates and adapts to the loss (*3*). Most people will also eventually form a new pair bond, which provides a behavioral indicator of loss adaptation (*4*). The processes that enable such adaptation remain poorly understood but likely occur via the same neural systems that are uniquely engaged by pair bonding (*5*).

Socially monogamous prairie voles are a laboratory-amenable species that recapitulates many aspects of human social bonds, making them ideal for interrogating the neurobiology of bonding and loss. Pair bonding in this species results in shift in sociobehavioral state where specific behaviors are only evident after the transition from living in a same-sex pair (i.e. family group) to an opposite-sex bonded pair. Pair bonded voles will prefer to affiliate with their partner, display selective aggression towards non-partner individuals, and exhibit robust and organized biparental care (*6–11*). Considering that other stable behavioral shifts, such as reproductive status and dominance hierarchies, are underwritten by changes in transcription, we aimed to determine if a mature pair bond—and the process of adapting to loss—are underwritten by stable changes in gene expression in the nucleus accumbens.

While an extended network of brain regions mediates social processing and decision making, the nucleus accumbens (NAc)—a region important for reward, motivation, and action selection—is a critical hub that is engaged when forming a bond and is implicated in loss processing (*12–14*). In humans, holding hands with a romantic partner enhanced blood oxygenation levels (BOLD) in the NAc and successful adaptation to spousal loss is associated with a reduced partner-associated BOLD signal in this region (*15, 16*). Thus, the NAc contributes to pair bonding and loss and broad scale changes in this brain region potentially mediate shifts in transcription which may be required to adapt to partner loss. Similarly, in prairie voles, pair bond associated behavioral changes are underwritten by several known neuromolecular changes within the NAc that maintain and reinforce pair bonds over time (*9, 12, 17*). Further, when separated from their partner, prairie voles exhibit behavioral and physiological distress that mirrors what is seen upon bond disruption in humans and other species, such as titi monkeys (*10*). These include increased circulating glucocortioids levels, activation of the HPA axis, increased anxiety, decreased pain thresholds, and autonomic dysfunction (*18–22*). However, we have previously shown that if a pair bonded prairie vole loses their partner, given enough time, they will form a new bond that supersedes the original bond (*23, 24*). As in humans, the ability to form a new bond indicates that prairie voles can adapt to partner loss.

Here we map the trajectory of the pair bond transcriptional profile by comparing opposite-sex paired males to same-sex cohoused naïve males. In the wild, sexually naïve male voles can cohabitate with other males, especially siblings, although these relationships do not persist after males form an opposite-sex pair bond and establish a nest/territory with their partner (*7, 10*). By comparing opposite-sex pair bonded voles to their same-sex paired, unbonded counterparts before and after partner separation, we have a ethologically relevant means to isolate the unique biology of bonding and loss independent of those that support general affiliative interactions (e.g. peer relationships) or the stressful effects of social isolation more broadly. Thus we compared behavior and NAc transcriptional profiles of opposite-sex pair bonded and same-sex housed naïve male voles to define the pair bond both behaviorally and transcriptionally. Then, we examined how the pair bond was altered specifically by partner separation to test the hypothesis that that pair bond transcriptional signatures and bond-associated behaviors would erode as a function of time since partner separation. We reasoned that these changes represent key components of loss adaptation that, together, may prime the vole to be able to form a new bond.

We found that pairing induced a reliable affiliative preference for a peer or a pair bonded partner, and that this preference is remarkably stable, persisting even after four weeks of separation. We further show that pair bond associated changes in accumbal gene expression remain consistent from two to six weeks post-pairing. However, once opposite-sex pairs are separated, the pair bond transcriptional signature erodes as a function of separation time. To further home in on the transcriptional changes associated with partner separation specifically in NAc neurons, we pioneered translating ribosomal affinity purification in voles (vTRAP). Using vTRAP, we identified clusters of genes associated with dopaminergic signaling, mitochondrial organization, and steroid hormone signaling whose expression patterns are sensitive to acute pair bond disruption and loss adaptation. In sum, our behavioral and transcriptional data suggests that erosion of pair bond transcriptional signatures in the NAc precedes changes in affiliative partner preference and selective aggression, providing insight into time-dependent neuromolecular changes that may contribute to loss adaptation.

## Results

We determined how bonding and extended separation affects social behavior and NAc transcription in opposite-sex and same-sex paired males. We employed timepoints that are experimentally validated and ethologically relevant (Fig. 1A). We began by pairing all study animals for 2 weeks, a duration that reliably produces mature pair bonds (*25, 26*). For the separation timepoints, the ability to form a new bond after loss serves as a behavioral metric of loss adaptation. Prior work has shown that male voles are able to form a new pair bond four weeks post-separation, but not two weeks or earlier (*23*).

**Figure 1.**
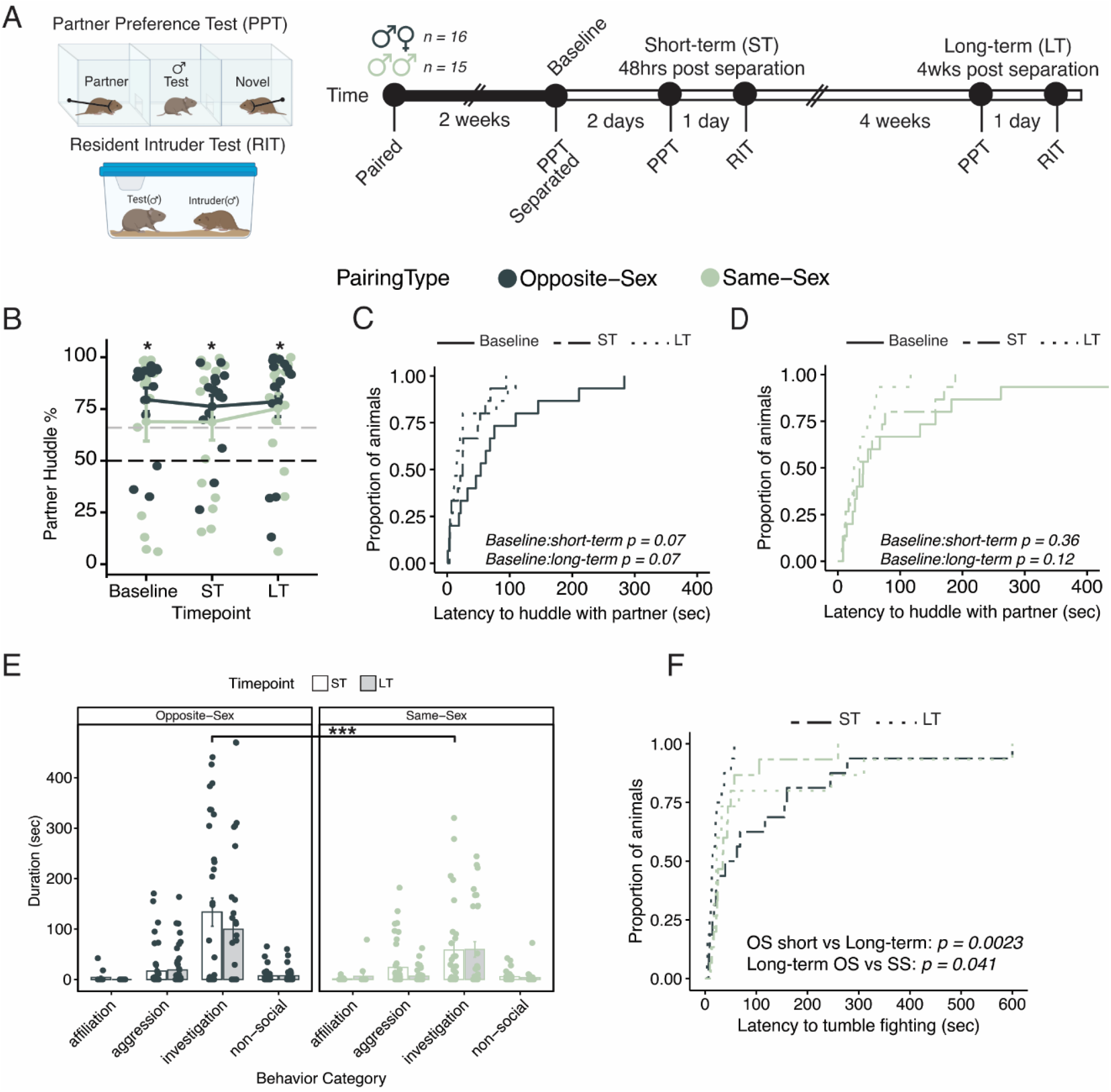
Males in opposite-sex pairs retain their partner preference and selective aggression for at least four weeks following partner separation. **(A)** Schematics of behavioral tests and timeline for behavioral experiments. Opposite-sex (OS; n = 16) and same-sex (SS; n = 15) pairs were paired for 2 weeks prior to a baseline partner preference test (PPT). Pairs were then separated for 48 hrs prior to short-term (ST) behavior tests of PPT and resident intruder (RIT) administered 24 hrs apart. Animals remained separated until the same behavior tests were repeated 4 wks later for the long-term (LT) time point. **(B)** Partner preference scores (% partner huddle/total huddle) from baseline, short-term, and long-term PPTs of opposite-sex (dark green) and same-sex (light green) paired males. Opposite- and same-sex paired males showed a baseline partner preference (one way t-test relative to 50%; opposite-sex: T_15_ = 5.14, p < 0.001, same-sex: T_15_ = 2.014, p = 0.064) that remained evident for both groups after short-term separation (opposite-sex: T_15_ = 3.908, p < 0.001, same-sex: T_15_ = 2.149, p = 0.050) and long-term separation (opposite-sex: T_15_ = 3.908, p < 0.002, same-sex: T_15_ = 3.477, p = 0.004). Black dotted line at 50% indicates no preference and gray dotted line indicates 2/3 of time with partner. **(C, D)** Cumulative event plots indicating that opposite-sex males huddle with their partner more quickly after separation (C: log-rank test: χ^2^ =5.6, p = 0.06) while same-sex males show minimal changes in partner huddle latency (D: log-rank test: χ^2^ = 4.9, p = 0.087). **(E)** Bar graph of the duration (sec) of each behavior in the resident-intruder test for opposite-sex (left) and same-sex (right) after short-term (white) and long-term (grey) separation. There is a significant difference in social investigation between opposite- and same-sex paired males at the short-term time point (3-way RM-ANOVA with Tukey’s post-hoc: p = 0.00097) but no other differences were observed. **(F)** Cumulative event plot showing latency to tumble fighting (sec) in the resident intruder test. After long-term separation, opposite-sex males were significantly faster to engage in tumble fighting, a marker of intense aggression, compared to the short-term time point (log-rank test: χ^2^= 9.30, p = 0.0023) or when compared to same-sex paired males (log-rank test: χ^2^= 4.20, p = 0.041).

Consistent with our laboratory observations, wild male voles who lose a partner will remain at the nest for ∼17 days (*24*). Thus, we assessed behavior and transcription in paired voles two days and four weeks after separation— pre- and post-loss adaptation, respectively.

### Longitudinal assessment of pair bonding behavior before and after separation

Selective affiliation and aggression both reinforce pair bonds, and the ability to form a new bond likely relies on the erosion of one or both of these behaviors. We measured partner-directed affiliation via a partner preference test, which measures the amount of time spent huddling either with the partner or a novel vole tethered at opposite ends of an arena. To measure selective aggression, we used the widely-deployed resident intruder test. All sample sizes and comprehensive statistical results, including effect size estimates, are reported in **Supplementary Table 1**.

### Extended separation does not reduce partner preference

We hypothesized that opposite-sex pairs would have a partner preference at baseline and after short-term separation, when the pair bond is still intact, but not after long-term separation when the pair bond is ostensibly weakened (*23, 27*). Similarly, we anticipated that same-sex pairs would have a partner preference at baseline but would exhibit a faster and more robust loss of partner preference when separated. Both opposite-sex and same-sex pairs form a selective preference for their partner after 2 weeks of cohabitation (one sample t-test relative to 50%: opposite-sex p = 0.00015, same-sex p = 0.064) and retain their partner preference after short-term and long-term separation (one sample t-test relative to 50%: opposite-sex short-term p = 0.00023, opposite-sex long-term p = 0.0016; same-sex short-term p = 0.050, same-sex long-term p = 0.0037). Additionally, there was no significant difference in partner preference score between opposite-sex and same-sex pairs over time (**Fig. 1B**; 2-way RM-ANOVA: Main effect of partner (opposite-vs same-sex): F_(1,84)_ = 1.40, p = 0.24, η = 0.016; Main effect of time: F_(2,84)_ = 0.18, p = 0.83, η = 0.0043; Partner X time: F_(2, 84)_ = 0.12, p = 0.89, η = 0.0028). Other commonly used metrics, including time spent huddling and distance from either conspecific, also did not show significant differences between groups, over time, or as an interaction of these variables (**Fig. S1A-C**; **Table S1**). Together, these results indicate that prairie voles are capable of forming an affiliative preference for an opposite- or same-sex partner that remains intact despite prolonged separation.

### Subtle differences in opposite-sex and same-sex partner preference are evident in locomotion, latency to huddle, and behavioral consistency across tests

As none of the typically employed partner preference test metrics indicated pairing-type-dependent differences, we investigated more subtle behavioral phenotypes. We first asked if there were differences in locomotion or partner huddle latency to query if opposite- or same-sex paired animals spent more time investigating their social environment. We reasoned that opposite-sex-paired animals would be more vigilant in assessing novel animals and would be more motivated to interact with their partner than same-sex-paired animals. Opposite-sex-separated males had significantly greater locomotion at baseline and after short-term separation but were indistinguishable from same-sex paired voles after long-term separation (**Fig. S1D**, 2-way RM-ANOVA with Tukey’s post hoc: Baseline opposite-vs same-sex p = 0.0017, short-term opposite-vs same-sex p = 0.024, long-term opposite-vs same-sex p = 0.36). This increased locomotion may reflect an attempt to rapidly alleviate the stress induced by partner separation or increased vigilance in pair bonded males in a new environment (*28*). Males in opposite-sex pairs also began huddling with their partner faster over the course of separation while there were no differences in latency to huddle with a novel animal in either opposite-sex or same-sex male pairs **(Fig. 1C, D**; Table S1, log-rank test main effect of time points, trends reported for all p < 0.10: opposite-sex partner χ^2^ = 5.6, p = 0.06, novel χ^2^ = 0.60, p = 0.72; same-sex partner χ^2^= 4.90, p = 0.087, novel χ^2^= 1.30, p = 0.52; interaction time X partner (opposite-v same-sex) χ^2^= 13.90, p = 0.02; interaction time X novel (opposite-v same-sex) χ^2^= 2.40, p = 0.80). The decrease in huddle latency in opposite-sex paired males may reflect enhanced motivation to be with their absent partner.

We next examined behavioral consistency across partner preference tests. First, we examined the consistency in partner preference scores between time points for each pairing type. The partner preference change score was calculated by subtracting the percent partner preference between time points (short-term separation - baseline; long-term separation - baseline) (Fig. S1E). Following short-term separation, opposite-sex-separated animals showed more consistent partner preference scores than same-sex animals while long-term separation caused the opposite-sex and same-sex groups to be indistinguishable from each other (**Fig. S1E**; F-test of equality of variances: short-term:Baseline F_(14)_ = 0.21, p = 0.0054; long-term:Baseline F_(14)_ = 0.65, p = 0.43). Second, we examined the consistency of pair bond behaviors throughout separation by assessing the strength of correlations between individual behaviors from baseline to short-term and baseline to long-term in opposite-sex or same-sex separated animals (partner and novel huddle times, total huddle time, partner preference score, and locomotion) **(Fig. S1F**). Strong social behavioral correlations were observed in opposite-sex paired animals between baseline and short-term separation but not after long-term separation, suggesting that pair bond behaviors destabilize following extended separation.

Same-sex paired animals showed uniformly weaker correlations across all timepoints. However, locomotor behavior remained correlated across all timepoints for opposite- and same-sex voles, indicating long-term separation selectively destabilizes social behaviors.

### Long-term separation reduces selective aggression in same-sex but not opposite-sex paired voles

As has been reported previously, both opposite- and same-sex-paired males demonstrated aggression towards a novel male vole in a resident intruder test (*29*). We hypothesized that opposite-sex-paired animals would be aggressive after short-term separation, when the pair bond is still intact, but not after long-term separation, when the pair bond is weakened. In same-sex animals, we anticipated reduced aggression after long-term separation and overall less aggression than opposite-sex-paired males. We found that aggressive behavior did not differ between pairing types after short-term separation (2-way RM-ANOVA in **Table S1**). Only same-sex-separated animals showed a reduction in tumble fighting duration and total aggressive behaviors (summed duration of tumble fighting, chasing and defensive postures) between short-term and long-term separation (**Fig. 1E, Fig. S1G** stats **Table S1**). We also examined the latency to begin tumble fighting. Opposite-sex males initiated tumble fighting more quickly after long-term than short-term separation and, at the long-term separation time point, were faster to aggress than same-sex paired males (**Fig. 1F**, log-rank test: opposite-sex latency (short-vs long-term) χ^2^= 9.30, p = 0.0023; (long-term opposite-vs same-sex) χ^2^= 4.20, p = 0.04). Affiliative behaviors (huddling) and non-social behaviors (autogrooming and digging) did not differ significantly either by partner type or separation time (**Fig. 1E, Table S1**).

### The pair bond transcriptional signature is stable over time

Changes in many behavioral states, such as a shift in dominance or reproductive status, are supported by stable changes in transcription, although this has not been examined in the context of pair bonds (*30–32*). While prior work has demonstrated that mating and cohabitation in prairie voles result in transcriptional changes within the NAc, the consistency of these changes as long as the bond remains intact has yet to be assessed (*33–37*). Thus, we compared NAc transcription in opposite-versus same-sex-paired voles following either 2 or 6 weeks of pairing/cohabitation (**Fig. 2B**). By comparing opposite-sex to same-sex-paired animals, we identified transcripts specific to pair bonds compared with those associated with affiliative behavior more generally. For consistency, we limited transcriptional assessment to voles that had a baseline partner preference >50% (**Fig. 2B; Table S1**) (OS n = 15; SS n = 11).

**Figure 2.**
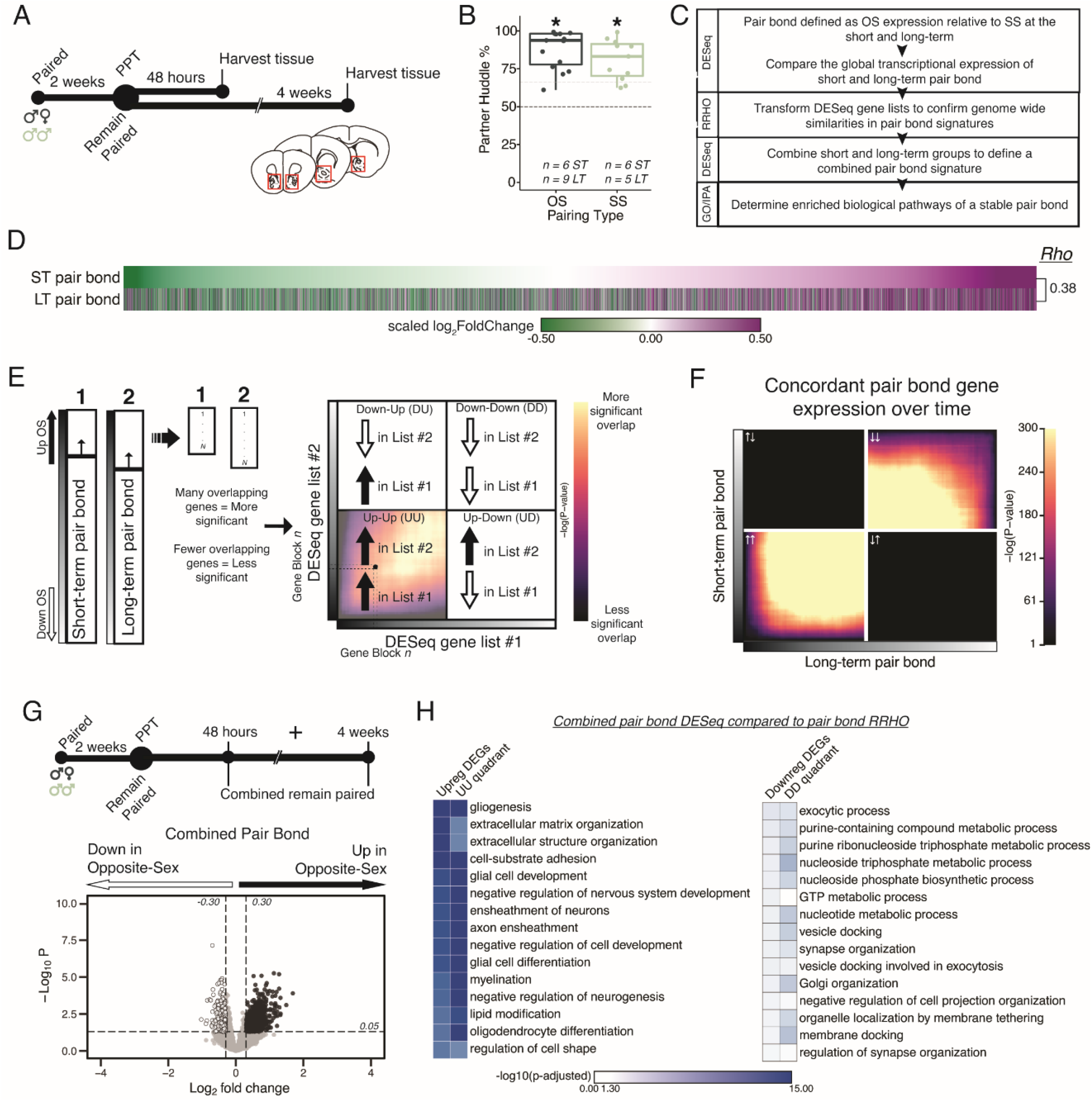
Pair bonding leads to persistent and consistent changes in NAc transcription. **(A)** Opposite- and same-sex pairs were paired for 2 weeks prior to a baseline partner preference test. Pairs then remain paired for either 48 hours (short-term; ∼2 weeks total pairing) or 4 weeks (long-term; ∼6 weeks total pairing) prior to collecting fresh nucleus accumbens tissue (dissection sites in red boxes) for RNA sequencing. **(B)** Baseline partner preference scores of males included in RNA sequencing for the opposite- and same-sex groups (one-tailed t-test relative to 50%: opposite-sex T_14_ = 11.76, p = 1.21 × 10^−8^; same-sex T_10_ = 7.78, p = 1.50 × 10^−5^). Black dotted line indicates a 50% partner preference score and the grey dotted line indicates 66%. There were no differences in partner preference score between opposite- and same-sex paired animals used for RNAseq (two-tailed t-test: T_21.016_ = 1.374, p = 0.184). **(C)** Transcriptional analysis workflow. **(D)** Gene list from both timepoints ordered from the smallest to largest log_2_FoldChange after short-term pairing with color indicating up- or down-regulation on opposite-vs same-sex pairs. Expression patterns are strongly correlated across timepoints (Rho = 0.38, p = 2.2 × 10^−16^). **(E)** Schematic of RRHO analysis. The heatmap is arranged into quadrants of genes upregulated in both lists (up-up: quadrant UU), downregulated in both lists (down-down: quadrant DD), or genes that have opposite regulation (up in list 1-down in list 2: quadrant UD; down in list 1-up in list 2: quadrant DU). Genes that are found in both lists at a similar ranked position result in higher p-values and are represented by a yellow color. **(F)** RRHO comparing short-term and long-term pair bonding (*from 2D*) indicates a stable pair bond gene signature over time as evidenced by concordant up- or downregulated genes at the two timepoints. **(G)** The short-term and long-term time points were pooled for opposite- and same-sex pairs to define the combined pair bond gene signature. **(H)** We compared GO analysis *Mus musculus* ontology terms between the combined pair bond DEGs and the RRHO quadrants (*from 2F*). Strong correspondence in the GO term significance between the two analyses supports that the combined pair bond transcriptional signature retains relevant biological information.

We used DESeq2 to identify transcripts up- or down-regulated in opposite-relative to same-sex paired males after short-term and long-term pairing (**Fig. S4A,B**) (*38*). When ordering transcripts based on their log_2_FoldChange at the short-term timepoint, the global transcriptional differences for opposite-versus same-sex pairs were strikingly similar following either 2 or 6 weeks of pairing, suggesting stable pair bond associated transcription across these timepoints (**Fig. 2D;** Spearman’s Rho = 0.38). We further determined that the observed correlation across timepoints is stronger than what would be expected by chance. We shuffled each animal’s cohort identity at the long-term timepoint and calculated Rho values between the observed short-term pair bond and the shuffled long-term pair bond over 1000 iterations (**Fig. S2A-D**). The cohort identity is a combination of three variables: pairing type (opposite-sex or same-sex), separation condition (remain paired or separated), and timepoint (short- or long-term). For example, long-term separated opposite-sex animals is one cohort identity of 8 total. This approach effectively randomized timepoint, partner type, and pairing status without disrupting underlying structure that exists within our transcriptional dataset due to correlations in expression across genes. Our observed Rho value was greater than 100% of iterations, indicating that the similarity in gene expression across short and long-term pairing timepoints is greater than would be expected by chance (**Fig. S2D**). Next, we asked whether our observed association across timepoints was driven by moderately but consistently expressed genes. Eliminating transcripts whose expression difference fell within the middle two quartiles in the short-term group resulted in a greater Rho value, indicating that the correlation across short- and long-term timepoints is more strongly driven by transcripts with larger expression differences between opposite- and same-sex pairs. To further interrogate biological function of altered transcripts, we set nominal thresholds of log_2_FoldChange > 0.30 or < -0.30, which are sufficient to identify replicable differences detectable via alternate methods (*39*), along with a p-value threshold of p < 0.05 to identify up- and down-regulated gene lists (DEGs). This less-stringent p-value threshold was used to generate robust enough gene lists to identify molecular pathways responsive to bonding with sufficient confidence while balancing the prohibitive costs associated with performing sequencing on 8 groups of animals. Prior studies in animal brain tissue using this nominal p-value have yielded biologically meaningful insights (*39–46*) (DEGs based on these thresholds are listed in **Table S2)**. There was significant overlap in the upregulated (104 shared, Fisher’s Exact Test: χ^2^ = 30.24, p = 5.36 × 10^−31^) and downregulated (46 shared, Fisher’s Exact Test: χ^2^ = 13.32, p = 7.05 × 10^−14^) transcripts across timepoints (**Fig. S4E**) (*47*).

To further examine similarity in transcriptional patterns between short-term and long-term pair bonds we employed Rank-Rank Hypergeometric Overlap (RRHO) (**Fig. 2E**) (*48, 49*). This threshold free approach allows us to determine if similar global transcription patterns are observed between two comparisons. The resulting RRHO heatmap is arranged into quadrants based on the direction of gene expression and each point represents the significance derived from the number of overlapping genes via the hypergeometric distribution (**Fig. 2E**). We determined how similar global transcription is between short-term and long-term pair bonds by comparing differential gene expression in the opposite-vs same-sex 2-week paired to opposite-sex vs same-sex 6-week paired groups. We observed extensive concordance of transcriptional patterns between 2- and 6-week paired voles, further supporting that the transcriptional profile of pair bonds is stable (**Fig. 2F**). Confirming that our observed RRHO signal was unlikely to be attributable to chance, we randomly shuffled cohort identity (**Fig. S2E**) and, separately, shuffled the gene ranking of the combined pair bond genes (**Fig. S6G**), both of which ablated the RRHO signal. Since the pair bond transcriptional profiles are largely stable over time, the ∼2 week and 6 week animals from each pairing type were pooled to create a single, well-powered opposite-sex-vs same-sex-paired comparison that defines the combined pair bond transcriptional signature (**Fig. 2G**). We then determined how this pair bond transcriptional signature changed following partner separation.

To determine which biological pathways might underlie the pair bond we employed Gene Ontology (GO) and Ingenuity Pathway Analysis (IPA) (*50*). Gene lists for GO terms or IPA were generated either from DEG lists or from genes in concordant (UU or DD) RRHO quadrants. GO terms that passed a false discovery rate of p < 0.05 were retained while IPA terms that had activation scores of at least -2 or 2 and whose enrichment for genes predicted to be regulated was significant (p-value < 0.05) were kept (**Supplemental Table 3**). Combined pair bond GO terms associated with the upregulated DEGs and the quadrant UU genes strongly implicate changes in glial cells and extracellular matrix organization (**Fig. 2H**), which is mirrored by activation of glioblastoma signaling via IPA (**Fig. S7A**). IPA additionally identified upregulation of Synaptogenesis, CREB signaling, and Endocannabinoid Neuronal Synapse pathways. All of these pathways are consistent with pair bonding as a form of complex learning that is mediated by neuromodulatory signaling (*14, 51–53*). Predicted upstream regulators of these pathways broadly support a role for learning and neuromodulation, including Creb1, Esr1, and various growth-factor and developmental genes (**Fig. S7A**). Combined pair bond GO terms associated with the downregulated DEGs the quadrant DD genes are implicated in nucleoside synthesis and synaptic plasticity (**Fig. 2H**). IPA indicates a suppression of corticotropin releasing hormone signaling, potentially reflecting the strong social buffering of stressors that occurs in pair bonded voles specifically (*20, 28, 54–56*) (**Fig. S7A**). Finally, the strong correspondence in the significance of GO terms between the combined pair bond DEGs and the RRHO quadrants further validates that pooling the time points retains biologically relevant transcriptional signatures (**Fig. 2H**).

### The pair bond transcriptional signature erodes following prolonged partner separation

We next tested the hypothesis that the pair bond transcriptional signature erodes following long-term partner separation. We paired opposite-sex and same-sex pairs for 2 weeks prior to a baseline partner preference test and then immediately separated males into new cages where they were singly housed for either 48 hours (short-term) or 4 weeks (long-term) prior to harvesting NAc tissue for RNAseq (**Fig. 3A**). As before, we only included animals with a baseline partner preference >50% (**Fig. 3B**). We determined the effect of separation on the pair bond gene signature by first defining differential gene expression of opposite-vs same-sex animals after short- and long-term separation and then used RRHO to examine genome-wide expression changes relative to the combined pair bond signature (**Fig. 3C**).

**Figure 3.**
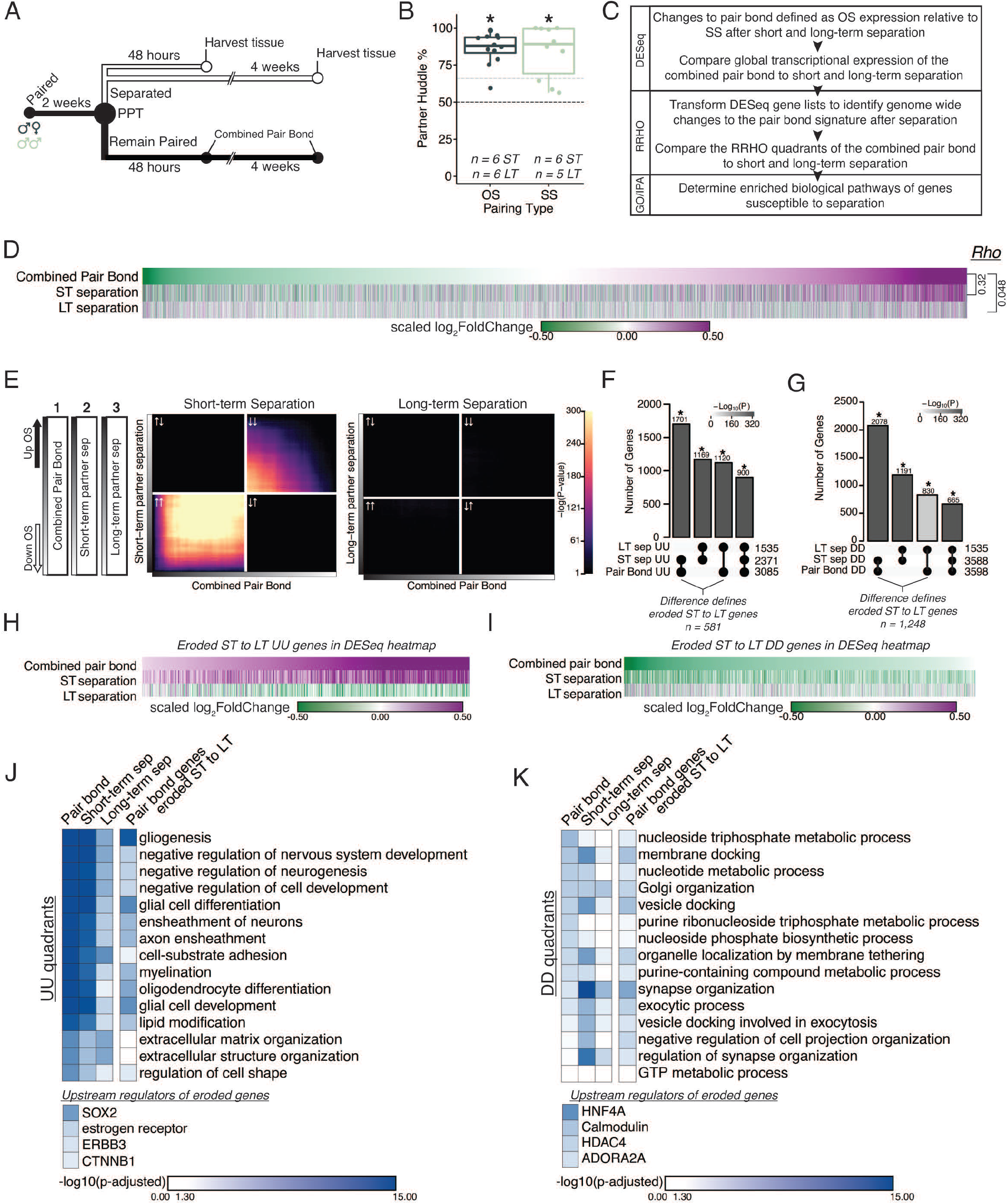
Prolonged separation erodes pair bond transcriptional signatures. **(A)** Opposite-sex and same-sex pairs were paired for 2 weeks prior to a baseline partner preference test. Pairs were then separated for either 48 hours (short-term) or 4 weeks (long-term) prior to collecting fresh nucleus accumbens tissue for RNAseq. Animals in the Remain Paired group are analyzed in Figure 2 to define the Combined Pair Bond transcriptional signature. **(B)** Baseline partner preference scores of males included in RNA sequencing for the opposite- and same-sex separation groups (one-tailed t-test relative to 50%: opposite-sex T_11_ = 11.56, p = 1.71 × 10^−7^; same-sex T_9_ = 6.07, p = 1.87 × 10^−4^). Black dotted line indicates a 50% partner preference score and the grey dotted line indicates 66%. There were no differences in partner preference score between opposite-sex and same-sex separated animals used for RNAseq (two-tailed t-test: T_14.33_ = 0.33, p = 0.75). **(C)** Transcriptional analysis workflow similar to Figure 2. **(D)** Heatmap of the scaled log_2_FoldChange for every gene where short- and long-term separation is compared to the combined pair bond gene signature. **(E)** RRHO using the combined pair bond, short-term separated, and long-term separated ranked transcript lists show that short-term separated animals retain a gene expression pattern concordant with pair bond transcription while there is a dramatic erosion of pair bond gene expression following long-term separation. **(F, G)** Upset plot showing overlap of genes in the UU or DD quadrants of *2F* and *3E* RRHO was determined using a Fisher’s Exact test. Mirroring the results from DESeq, more overlap is found between the pair bond and short-term separated gene lists than with long-term separated gene lists. The difference between the intersections of the pair-bond:short-term sep. and pair-bond:long-term sep. was used to define the eroded gene lists. **(H, I)** The RRHO gene lists of eroded pair bond genes were used to filter the DESeq heatmap in *2D*. Eroded genes from the UU quadrants show upregulation during pair bonding and short-term separation but downregulation after long-term separation **(H)** while the eroded genes from the DD quadrants show the opposite pattern **(I). (J, K)** GO and IPA analysis of RRHO quadrant gene lists and pair bond eroded gene lists. Scale represents the –log_10_(p-adjusted) where any non-significant terms are white. **(J)** Glial associated GO terms of the UU quadrants are less significantly represented after separation and are highly significantly represented among the eroded genes. Significant upstream regulators of the eroded genes include Sox2, estrogen receptor, Erbb3, and Ctnnb1. **(K)** Vesicle docking and synapse organization associated GO terms of the DD quadrants are less significantly represented after separation and are more significantly represented among the pair bond eroded genes. Significant upstream regulators of the eroded genes include Hnf4a, Calmodulin, Hdac4, and Adora2a.

First, we performed differential expression analysis of opposite-vs same-sex-separated males at each separation time point (**Fig. S4C,D**). There were significantly more shared DEGs between the combined pair bond group and short-term separation group than would be expected by chance, but this was not observed when comparing the shared genes between the combined pair bond and long-term separation group (**Fig. S4F**, Fisher’s exact test: pair bond:short-term sep χ^2^= 31.86, p = 6.78 × 10^−19^; pair bond:long-term sep χ^2^= 17.96, p = 0.53). Further, there are even fewer shared DEGs between short- and long-term separation suggesting notable transcriptional differences between these time points (**Fig. S4F**). The difference in transcription between short- and long-term separation is particularly evident when comparing the global transcriptomes through the lens of pair bond transcription. Specifically, when genes are ordered based on their degree of down- or up-regulation in the pair bond group we found that the combined pair bond signature remains intact after short-term separation (Rho = 0.32) but is largely eroded after long-term separation (Rho = 0.048) (**Fig. 3D**). We further assessed whether these correlations differed from chance via the previously employed permutation analysis. We found that the pair bond:short-term Rho value indicated a stronger association than would be expected by chance (p = 0.0362) while the pair bond:long term separation association value was highly likely to occur by chance (p = 0.41). Together, these results suggest that extended separation erodes nucleus accumbens pair bond transcription, a potentially crucial molecular process needed to prime the vole to be able to form a new bond.

To more comprehensively assess genome-wide changes to the pair bond signature following extended partner separation we again employed RRHO (**Fig. 3E**). We compared the ranked gene list of opposite-vs same-sex for the 1) combined pair bond gene signature (**Fig. 3D, top**) versus either 2) short-term partner separation (**Fig. 3D, middle**) or 3) long-term partner separation (**Fig. 3D, bottom**). We found that the pair bond signature remains largely intact after short-term separation as indicated by significant signal among upregulated genes (quadrant UU), and to a lesser extent, downregulated genes (quadrant DD) (**Fig. 3E**). However, following long-term separation, the pair bond gene signature is largely undetectable as indicated by a lack of significant signal throughout the RRHO plot. Plotting the long-term RRHO with an unadjusted p-value scale revealed few overlapping genes in the UU and DD quadrants. The low level of overlap indicates that there is a residual, albeit greatly reduced, pair bond signature that is still detectable after long-term separation (**Fig. S6C**). Additionally, we used the same control analyses as previously employed (**Fig. S6G-I**).

We next examined which genes are consistently present in the UU or DD quadrants between pair bonding and each separation timepoint. Specifically, we compared the UU and DD quadrants (inclusive of all transcripts in each quadrant) of the short-term (2 week) vs long-term (6 week) pair bond from Fig. 2F, the combined pair bond vs short-term separation in the left side of Fig. 3E, and the combined pair bond vs long-term separation on the right side of Fig. 3E. The degree of overlap in the UU quadrants was greatest between the combined pair bond vs short-term separation and least between the combined pair bond vs long-term separation, a result consistent with the most extensive erosion of pair bond transcription occurring after long-term separation (**Fig. 3F**: all stats in **Table S1**, genes in **Table S2**). The same trend was observed for transcripts in the DD quadrants (**Fig. 3G**, all stats in **Table S1**, genes in **Table S2**). We next identified genes whose differential expression in pair bonded voles eroded as a function of separation time. We reasoned that such genes are involved in biological pathways that likely regulate adapting to partner loss and prime an animal to form a new pair bond. The genes that were no longer present in the pair bond:long-term separation intersection that were present in the pair bond:short-term separation intersection were deemed eroded genes. Pair bond transcripts no longer found between short-term and long-term separation (i.e. eroded transcripts) include 581 UU quadrant transcripts and 1,248 DD quadrant transcripts (**Fig. 3F, G**). We next filtered the genome-wide expression data of Fig. 3F by the eroded gene lists (**Fig. 3H, I**). The eroded UU genes are strongly upregulated in a stable pair bond and show dramatic downregulation following long-term separation (**Fig. 3H**). Thus, the eroded UU genes and their associated biological processes may represent key aspects of adapting to partner loss. There was also a corresponding reduction in DD gene overlap between a stable pair bond and following separation, further supporting that bond-related changes in gene expression erode after long-term separation regardless the initial direction of expression during bonding (**Fig. 3I**).

Finally, we determined how the separation induced transcriptional changes affected the underlying pair bond associated biological processes. We used the pair bond associated GO terms from Fig. 2H as a reference point to determine if these terms were still significantly associated with the underlying transcriptional profile following partner separation (**Fig. 3J, K**). We predicated two outcomes. First, separation would reduce the significance of the pair bond associated GO terms and second, pair bond genes whose expression eroded would be significantly represented by the original pair bond associated terms. The transcripts in the short-term separation UU quadrant were still significantly associated, though to a lesser degree, with the pair bond associated GO terms. After long-term separation the underlying transcriptional signature is much less significantly associated with the pair bond GO terms. Further, the eroded transcripts are highly associated with the pair bond associated GO terms, including glial-associated terms. A potential role for glia was also implicated in the IPA identification of pathways for glioblastoma multiforme signaling and glioma signaling (**Fig. S7C**). Upstream regulators inducing the erosion of these glial-associated genes include Sox2, estrogen receptor, Erbb3, and Ctnnb1 (**Fig. 3J, Fig. S7C**). We observe a similar trend in the DD quadrants where the pair bond associated GO terms become less representative of the underlying transcriptional signature following long-term separation but are associated with the eroded genes, though to a lesser degree (**Fig. 3K**). IPA pathways activated in a stable pair bond (endocannabinoid signaling, synaptogenesis, and dopamine signaling) are significantly represented among the eroded DD genes (**Fig. S7C**). Upstream regulators for the eroded DD transcripts include Hnf4a, Calmodulin, Hdac4, and Adora2a (**Fig. 3K, Fig. S7C**).

### Neuronal interrogation of long-term partner separation reveals gene clusters associated with pair bond disruption and loss adaptation

Our tissue-level analysis of gene expression strongly implicated a role for glia in bond formation and loss, and we reasoned that relevant neuronal transcriptional changes may have been masked by these prominent glial transcriptional signals. Thus, we specifically examined separation-induced neuronal transcriptional changes by pioneering translating ribosome affinity purification in voles (vTRAP) (*57, 58*). The vTRAP construct, DIO-eGFP-RPL10a, is a double-floxed inverse orientation eGFP-tagged ribosomal subunit that uses the prairie vole RPL10a gene sequence (**Fig. 4A**; **Fig. S8A**). The vTRAP construct is paired with Cre-recombinase for inducible, cell-type specific expression. AAV-mediated delivery of hSyn-Cre, which drives neuron-specific Cre-recombinase expression, and hSyn-DIO-vTRAP vectors results in neuronal expression of GFP-tagged ribosomes. Subsequent immunoprecipitation of the tagged ribosomes provides a means to isolate neuron-specific, actively-translating mRNAs (*57, 58*). vTRAP is Cre-dependent as confirmed by bilateral NAc injections of +/- Cre (**Fig. S9A**). We performed vTRAP immunoprecipitation of 3 cohorts: 1) opposite-sex long-term separated, 2) opposite-sex long-term remain paired, and 3) same-sex long-term separated.

**Figure 4.**
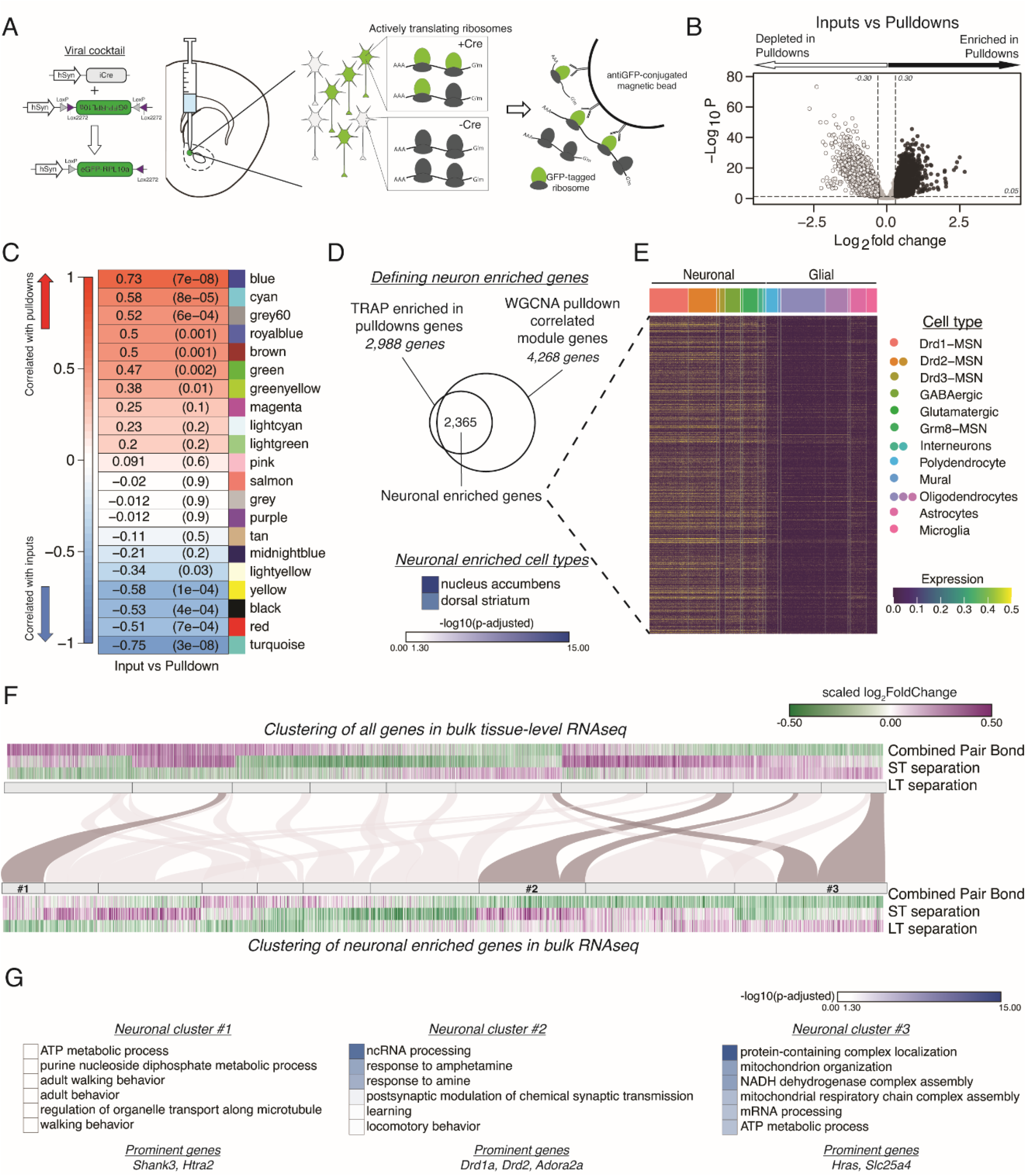
vTRAP elucidates separation-induced transcriptional changes specifically in neurons. **(A)** Schematic of the vTRAP system. AAV-mediated delivery of hSyn-Cre, which drives neuron-specific Cre-recombinase expression, and DIO-vTRAP vectors results in neuronal expression of GFP-tagged ribosomes. Subsequent immunoprecipitation of the tagged ribosomes isolates neuron-specific, actively-translating mRNAs. **(B)** Transcripts enriched in the immunoprecipitation pulldowns compared to the input fractions. Differential expression skews towards depletion. **(C)** WGCNA to identify gene modules correlated with the input (negative correlation) or pulldown (positive correlation) fractions. The grey60 module contains neuronal transcripts of interest such as Drd1a, Fosb, Oprk1, and Pdyn while the blue module contains Drd2. **(D)** Neuronal enriched genes—2,365 genes—were defined as the intersection of the Input v Pulldown enriched DEGs and the WGCNA pulldown-correlated modules (modules grey60, blue, greenyellow, brown, green, royalblue, and cyan). When using Enrichr to search for cell types associated with the neuronal enriched gene list, nucleus accumbens is the most prominent and is highly significant followed by dorsal striatum. Scale represents –log_10_(p-adjusted). **(E)** Heatmap showing expression of identified neuronally enriched genes in a single-nuclei RNAseq dataset from the nucleus accumbens of adult male rats. The scale indicates expression of each gene in the snRNAseq dataset. **(F)** Hierarchical clustering using one minus Pearson correlation with complete linkage of bulk tissue-level sequencing for all genes (top heatmap) and neuronal enriched genes (bottom heatmap) for the combined pair bond, short-term separation, and long-term separation. The scale indicates scaled log_2_FoldChange. The mapping of neuronal clusters to the all genes clusters is visualized via a Sankey plot. Neuronal clusters of interest are highlighted by dark grey links. **(G)** Gene Ontology analysis using mouse terms for each neuronal cluster of interest. Genes of interest that occur in multiple terms are referred to as prominent genes. Scale represents –log_10_(p-adjusted). GO terms associated with neuronal cluster #1 are not significant due to the small number of genes in that cluster.

We defined a set of neuronally enriched genes by using the intersection between threshold and threshold-free based analyses. We first identified neuronally-enriched transcripts by comparing the transcriptional profiles of the input fraction (equivalent to bulk RNA-seq) and pulldown fraction of the animals from the TRAP groups specified above. We predicted that the pulldown fraction would be enriched for neuronal markers and depleted in glial-associated transcripts. Differential expression analysis using the previously defined thresholds revealed that most transcripts are depleted in the pulldown fraction as expected (**Fig. 4B**). GO terms support that hSyn-vTRAP successfully isolates neuronally-enriched transcripts and filters out genes associated with gliogenesis (**Fig. S9C**). Second, we used weighted correlation network analysis (WGCNA) to identify gene modules significantly correlated with the input (negative correlation) or pulldown (positive correlation) fractions (**Fig. 4C, Fig. S5B**) (*59*). Of note, two of the most significantly positively correlated modules, blue and grey60, contain multiple neuronal genes of interest such as Drd1a, Drd2, Oprk1, Fosb, and Pdyn, which have previously been implicated in pair bonding and reward learning (*35, 36, 60*). Enrichr analysis for cell type also indicated that our identified neuronal enriched genes are predominantly associated with the nucleus accumbens and dorsal striatum (**Fig. 4D**) (*61–63*). Finally, we validated that the gene list at the intersection of the DEGs and the WGCNA modules was predominately expressed in neurons and not glia using a publically available single nuclei nucleus accumbens gene expression dataset generated from adult male rats (*64*). We filtered the dataset by our neuronal enriched gene list and plotted those gene’s expression in the rat nucleus accumbens which confirmed that our genes are predominately expressed in neuronal cell types (**Fig 4E**).

We next queried potential neuronal transcriptional changes in our tissue-level RNAseq data. To ensure that the tissue-level expression is representative of neuron-specific expression we compared the expression patterns of the neuronal enriched genes in the input and pulldown fractions of opposite-vs same-sex long-term separated males (**Fig. S9D**). The strong similarity in expression validates that bulk sequencing expression can be used to infer the neuronal expression of those same genes (**Fig. S9D**). As such, we filtered the tissue-level RNAseq dataset from Fig. 3D by the neuronal enriched gene list of Fig. 4D to track neuronal gene expression throughout partner loss. We then performed unsupervised clustering of all genes (**Fig. 4F**, top) and neuronal enriched genes (**Fig. 4F**, bottom). We identified three neuronal-enriched gene clusters that had particularly interesting expression patterns. Cluster 1 consists of genes that were upregulated in opposite-sex paired males during pair bonding and short-term separation and downregulated following long-term separation. GO terms associated with these genes include adult walking behavior, a term that contains the autism-associated gene Shank3 (**Fig. 4G**) (*65, 66*). Cluster 2 transcripts are downregulated in pair bonded males and then robustly upregulated in opposite-sex separated males at the short-term and, to a lesser extent, at the long-term timepoint indicating an acute transcriptional response to pair bond disruption. The GO terms associated with these genes suggest an involvement of dopaminergic systems (Drd1a and Drd2) and learning processes potentially engaged to adapt to loss (**Fig. 4G**). Finally, Cluster 3 contains genes that were downregulated in pair bonded males and became upregulated only after long-term opposite-sex partner separation, representing transcripts that may help prime the vole to form a new bond. The GO terms associated with this cluster are primarily implicated in cellular metabolism (**Fig. 4G**). IPA revealed that neuronal clusters 2 and 3 were also enriched for estrogen receptor signaling and oxidative phosphorylation pathways with cluster 2 also implicating opioid signaling pathways (**Fig. S7D**). Steroid and opioid signaling have been extensively implicated in pair bonding (*35, 36, 51, 67–71*).

## Discussion

Our data provide the first comprehensive assessment of social behavior and accumbal transcription in pair bonded male prairie voles before and after partner loss, providing novel insight into the biological processes that may enable loss adaptation. Prairie voles, unlike laboratory mice and rats, form exclusive lifelong pair bonds that are distinct from affiliative same-sex relationships, making them ideal for studying the unique neurobiology of bonding and loss (*7, 10, 21, 22, 29, 72, 73*). Additionally, the direct comparison of opposite-versus same-sex paired animals provides an important control for the effects of social context – whether living with another animal or alone (*74, 75*). By leveraging this comparison across our experiments, we demonstrated that pair bonding leads to consistent and persistent shifts in NAc transcription, which erode as a function of long-term partner separation. These changes in NAc transcription were accompanied by only subtle differences in behavior. This suggests that substantial changes in accumbal gene expression may contribute to priming the vole to form a new bond but are not sufficient to erase an existing bond. In addition, we demonstrate the utility of vTRAP for interrogating neuron-specific responses to bonding and separation that are obscured when analyzing bulk sequencing results. Together, these results expand the utility of monogamous prairie voles for future behavioral and molecular-genetic studies of partner loss and loss adaptation.

### Partner preference persists despite extended separation

We observed overall similarity and persistence in social behaviors exhibited by opposite-sex and same-sex paired prairie voles, both of which displayed a partner preference despite prolonged separation. The partner preference test, as employed in our study, may be insufficient to capture the uniquely rewarding aspect of pair bonds compared with other affiliative relationships. In part, this may reflect the relatively crude but commonly employed metrics used to infer partner preference, which rely on proximity to the tethered animals without analyzing the valence or content of specific social interactions. Despite the similarity of behavior between both pairing types, pair bonds are known to be more rewarding than same-sex affiliative relationships. Prairie voles will show a conditioned place preference for a context previously paired with their partner but not for a context paired with a same-sex cagemate (*72*). A more nuanced assessment of behavior would likely further elucidate the subtle differences in partner-directed behavior that we observed, such as latency to huddle with the partner and correlation across many partner preference behaviors.

While our results contradict prior work demonstrating dissolution of partner preference in male voles after four weeks of separation (*27*), this may be due to differences in study design. In particular, prior studies paired male and female voles for much shorter periods of cohabitation prior to separation and examined behavioral pair bond dissolution in different cohorts of animals at each timepoint (*10, 27*). We instead paired animals for two weeks prior to separation. Pair bonds strengthen and mature over this extended cohabitation period, which may contribute to the observed perseverance of partner preference for weeks after separation (*25, 26*). In addition, our longitudinal study design resulted in re-exposure to the partner in the partner preference test performed 48 hours post-separation. This brief reunion may serve to reinforce the bond in a way that does not occur if the test animal is not re-exposed to their partner post-separation. Finally, we employed tubal ligation, an approach that sterilizes females while leaving them hormonally intact. There is conflicting work suggesting that pregnancy can affect pair bonding behaviors while other studies find no difference in male partner preference between intact and ovariectomized females (*12, 26, 76, 77*). Regardless, our results suggest that male voles have the capacity to remember and potentially resume an affiliative relationship – pair bonded or not - despite prolonged separation and isolation.

### Pair bonding results in consistent glia-associated transcriptional changes that erode following prolonged separation

We first asked whether bonding-induced gene expression is stable as long as pairs remain cohoused (*35–37, 78*). Using three orthogonal analytical methods, we found remarkable consistency in the transcriptional pattern of opposite-vs same-sex pairs co-housed for 2 or 6 weeks. Prior work on social dominance suggests that such stable transcriptional changes are critical to maintain different behavioral states (*30–32*). However, to our knowledge, this is the first demonstration that a similar conceptual underpinning may maintain mature pair bonds.

Consistently upregulated transcripts in pair bonded males are associated with glia and extracellular matrix organization while downregulated transcripts are associated with nucleoside processes, synaptic plasticity, and vesicle trafficking. Together, these terms implicate processes central to neuronal remodeling and synaptic plasticity. Of note, while our analyses do identify a number of mitochondrial GO terms, these terms are not as strongly implicated in our analyses as in previous reports (*33*). Despite this, our study confirms a potential role for mitochondrial dynamics in the NAc in male prairie vole pair bonding. This is consistent with a model in which pair bonding and separation induce neuronal plasticity and synapse re-organization, which require large amounts of energy and induce mitochondrial redistribution within neurons (*79, 80*).

We next found that pair bond transcriptional signatures erode as a function of separation duration. Via RRHO, we found that there is a considerable reduction in concordant pair bond gene expression following long-term, but not short-term, separation. When we asked which transcripts and associated biological processes erode over this timeframe, we found that they largely corresponded with those evident in the stable pair bond transcriptional signature. Specifically, transcripts associated with glia, extracellular matrix organization, synapse organization, and vesicle docking are stably expressed in pair bonded males but eroded following long-term separation. The erosion of the pair bond transcriptional signature over time in opposite-sex separated males may be central to adapting to partner loss in order to form a new pair bond if given the opportunity.

Our analyses strongly indicate a role for glia in regulating pair bonding and the response to partner separation. Transcripts associated with gliogenesis, glial cell differentiation, and myelination are upregulated in stable pair bonds, potentially reflecting the importance of glial cells in sculpting and refining neural circuits in response to social experience (*51, 81, 82*). Similar GO terms—with the addition of oligodendrocyte differentiation—are associated with eroded pair bond transcripts after long-term partner separation. Hypomyelination and oligodendrocyte dysfunction are seen in mice and rats after chronic social stress and in individuals with depression (*83–85*). One mechanism for social deprivation-mediated hypomyelination and impaired oligodendrocyte maturation is by reduced flux through Erbb3-based signaling pathways (*86, 87*). Of note, Erbb3 is one of 24 genes whose differential expression is conserved in taxonomically diverse monogamous species when compared to closely-related non-monogamous relatives (*88*). We find Erbb3 upregulated during pair bonding through short-term separation but it is no longer upregulated following long-term separation. Further, IPA analysis indicates that Erbb3 is an upstream regulator of the eroded genes. Thus, it is possible that long-term separation is downregulating Erbb3-associated pathways resulting in reduced myelination and disrupting oligodendrocytes specifically in opposite-sex-separated animals. In adult mice, social reintroduction rescued hypomyelination suggesting a role for oligodendrocyte plasticity in social behavior (*87*). An important future direction is whether similar plasticity occurs in voles; does bonding with a new partner re-engage these glial processes?

Finally, we reasoned that the prominence of glial-related terms may have masked important neuronal transcriptional changes. To home in on neuronal transcriptional changes specifically, we developed vole-optimized translating ribosome affinity purification in prairie voles (vTRAP). We used DESeq, WGCNA, and a publicly available snRNAseq dataset to identify neuronal enriched genes, which show a 1.2 – 8.8 fold enrichment in neuronal pulldowns compared to the tissue-level sequencing data. We then queried expression changes specifically among these genes as a function of pair bonding and partner separation. With this enhanced sensitivity, we were able to identify three neuronal gene expression patterns that are differentially sensitive to partner loss. Notably, dopamine associated genes are robustly upregulated in neurons after acute bond disruption, possibly reflecting a withdrawal-like state (*89–91*). Genes associated with anxiety and energy production are upregulated only after long-term separation, possibly to facilitate finding a new partner. Together with our tissue level sequencing implicating the role of glia, vTRAP provides a more nuanced understanding of how the pair bond neuronal transcriptional signature responds to partner separation over time.

### Limitations and future directions

A notable aspect of our results is that the substantial erosion of pair bond transcription in the NAc after long-term separation is not matched by a similar magnitude change in behavior, despite evidence that changes in gene expression in this brain region help maintain bonds over time (*12, 37, 78*). This suggests that even long-term separated pairs retain a residual pair bond associated transcriptional signature that may be sufficient enough to influence behavior. One potential explanation for this result is that our study only examined responses in the NAc, and the neurogenomic signature of pair bonding may persist in other relevant brain region(s) (*13, 34, 55, 92*). The medial pre-frontal cortex and the hippocampus innervate the NAc and are implicated in pair bonding related behaviors and sociocognitive forms of memory, representing important targets for future research (*93–96*). Additionally, pair bonding modifies dopamine release into the NAc and this change in dopamine tone may be due to long-lasting changes in ventral tegmental area neurons (*35*). Another possible explanation is that the residual pair bond transcriptional signature may reflect a form of epigenetic memory, and/or long-lasting proteomic changes may be sufficient to maintain partner preference and selective aggression in opposite-sex-separated males (*97, 98*). Alternatively, transcriptional changes in a small subset of cells, which may be undetectable in tissue-level RNAseq, could contribute to the persistence of partner preference. To address the latter, we began to narrow our inquiry through the development of vTRAP to isolate actively translating mRNAs from specific cell types—in this case neurons. Recent advances in cell-type and projection-specific targeting via Cre recombinase and cell-type specific promoters deployable via AAV will further broaden application of vTRAP in voles (*99, 100*). Additionally, refined approaches, such as single-cell and single-nucleus RNAseq, have the possibility of greatly enhancing our understanding of the specific cell populations regulating complex behaviors.

Finally, it is worth noting that the persistence of partner preference in our paradigm may have been facilitated by the study design. Unlike in the wild where there are potential opportunities to rebond upon partner loss, the separated males in this study had no opportunity to form a new bond as they were singly-housed. Under such conditions, it may be advantageous to maintain behaviors that would revive the bond if a partner returned. A more ethologically relevant question may be how long it takes before a separated vole is capable of forming a new bond. Previous work from our lab has shown that male voles can form a new, stable pair bond that supplants the original bond after four weeks of separation but not sooner (*23*). This would suggest that although partner preference remained intact in the current study, these males were likely capable of forming a new bond if the opportunity arose, a capacity that may arise from the erosion of pair bond associated transcription. Such an explanation is parsimonious with the human experience. We do not forget our previous bonds; rather we integrate the loss in order to continue on with life.

In sum, we provide foundational assessments that are integral for establishing prairie voles as a relevant model for studying loss. Here we describe that, in the absence of new bonding opportunities, the original pair bond will persist for at least 4 weeks post-partner-separation. However, prior work indicates that there is still an adaptive process occurring within this time frame that primes a vole to form a new bond if available (*23*). Our work suggests that this adaptation is mediated at least in part by an erosion of bond-related transcription in a reward-related brain region that largely precedes bond dissolution at the behavioral level. Future studies are needed to ask whether manipulating the direction of key pair bond transcriptional changes is sufficient to facilitate re-bonding with a new partner. Together, this work and future studies may help shed light on the pain associated with loss, as well as how we integrate grief in order to meaningfully reengage with life.

## Materials and Methods

### Animals

Sexually naïve adult prairie voles (*Microtus ochrogaster*, n = 232: 172M, 60F, P60 - P168 at experiment start) were bred in-house. Prior to cohabitation, female partners were tubally ligated to control for the confounds of pregnancy and recovered for 2 weeks. Both opposite-sex and same-sex pairs cohabitated for 2 weeks prior to baseline partner preference tests. While mating was not confirmed, this duration of cohabitation results in pregnancy in at least 90% of intact pairs, and the pair bonds in intact animals assessed in a separate study in our lab were behaviorally indistinguishable from those in the current study (t-tests: partner preference score t_(28.7)_ = 0.39, p = 0.70; partner huddle duration t_(28.6)_ = -0.29, p = 0.78; novel huddle duration t_(24.5)_ = -0.81, p = 0.42; total distance traveled t_(26.3)_ = -1.86, p = 0.075) (*25, 26*). Interestingly, the presence of young at the nest also does not influence the amount of time a male remains after losing his partner in the wild (*24*). Separated pairs were distanced from each other in the colony room. All procedures were performed in accordance with standard ethical guidelines (National Institutes of Health Guide for the Care and Use of Laboratory Animals) and approved by the Institutional Animal Care and Use Committee at the University of Colorado Boulder.

### Behavioral assays

Partner preference tests (PPT) were performed as previously described (23) with post-hoc tracking using Topscan software (v3.0) and analysis using a custom Python script. Resident intruder tests (RIT) were performed in the experimental male’s home cage with an intruder male for 10 minutes and behaviors were hand scored by a trained observer blind to conditions using BORIS (v7.10.2) (24). All behaviors were performed between 09:00 and 14:00.

### RNA Sequencing

Nucleus accumbens dissection and RNA extraction of animals with a baseline partner preference was performed as described in Heiman *et al*. (25). Library preparation was via the KAPA mRNA HyperPrep kit with polyA enrichment for 1×75 RNA sequencing using Illumina NextSeq V2 for a total read depth of ∼20M reads/sample.

### Statistics and Bioinformatics

All sequencing analysis was performed using R (v4.0.4) (*101*). For transcriptomic analysis, reads were aligned to the Ensembl prairie vole genome 1.0 and only genes with >10 reads were retained. Pairwise differential expression comparisons were done using DESeq (v1.30.1) (26) using a threshold of log2FoldChange +/- 0.30 and p-value < 0.05. Threshold-free analysis of directional regulation was accomplished using RRHO2 (v1.0) (27, 28). Gene overlap and Gene Ontology overlap was done using the SuperExactTest package (v1.0.7) (29) and the clusterProfiler package (v3.18.1) (30). Ingenuity Pathway Analysis (Qiagen) was used to predict upstream regulators and molecular pathways as in (*39*). The R package Weighted Gene Correlation Network Analysis (WGCNA) (v1.70-3) only used the vTRAP cohorts to identify neuronal enriched genes. Plots were made using base plot, ggplot2 (v3.3.3), EnhancedVolcano (1.8.0), RRHO2 (v1.0), clusterProfiler (v3.18.1), and Adobe Illustrator (25.2.3) (*49, 50, 102, 103*).

## Author Contributions and Notes

Z.R.D. and J.M.S developed experimental design. J.M.S., C.J.K., X.G.B. executed experiments. J.M.S, L.E.B., D.M.W, and Z.R.D. analyzed and interpreted data. Z.R.D., L.E.B., and J.M.S. wrote and edited the manuscript.

The authors declare no conflict of interest.

All sequencing data will be assigned a GEO accession number by the time of publication. All code will be available on the Donaldson Lab GitHub (https://github.com/donaldsonlab) and all additional data will be available on Figshare. The vole optimized DIO-eGFP-RPL10a vector will be made available on Addgene.

## Acknowledgments

We thank Robin Dowell, Mary Allen, Ryan Logan, Xiangning Xue, and Gracie Sapp for their assistance in designing the experiments and analysis. We also thank Cayla Jo Paulson and Jessica Abazaris of the animal care staff at the University of Colorado Boulder. We thank the rest of the Donaldson lab for their feedback and support and the voles for their sacrifice. For vTRAP images we used the Molecular, Cellular, and Developmental Biology Light Microscopy Core at the University of Colorado Boulder with James Orth’s assistance. This work was supported by NIH award DP2OD026143 and funds from the Whitehall Foundation and the Dana Foundation (to Z.R.D.) and NIH award T32 GM008759-17/18 (to J.M.S.).

## Supplementary Information Text

### Materials and Methods

#### Animals

Sexually naive adult prairie voles (*Microtus ochrogaster*, N = 232: 172M, 60F, P60 - P168 at experiment start) were bred in-house in a colony originating from a cross between voles obtained from colonies at Emory University and University of California Davis, both of which were established from wild animals collected in Illinois. Animals were weaned at 21 days and housed in same-sex groups of 2 – 4 animals in standard static rodent cages (7.5 × 11.75 × 5 in.) with ad-lib water, rabbit chow (5326-3 by PMI Lab Diet) supplemented with alfalfa cubes, sunflower seeds, cotton nestlets, and igloos for enrichment until initiation of the experiment. All voles were aged between post-natal day 60 and ∼180 at the start of the experiment. Throughout the experiment, animals were housed in smaller static rodent cages (11.0 in. x 8.0 in. x 6.5 in.) with ad-lib water, rabbit chow (5326-3 by PMI Lab Diet), and cotton nestlets. They were kept at 23–26°C with a 10:14 dark:light cycle to facilitate breeding. All procedures were performed in accordance with standard ethical guidelines (National Institutes of Health Guide for the Care and Use of Laboratory Animals) and approved by the Institutional Animal Care and Use Committee at the University of Colorado Boulder.

#### Tubal ligations

All females in opposite-sex pairs were tubally ligated to avoid confounds of pregnancy, but to keep the ovaries intact as to not impact hormonal function. Females were anaesthetized with a mixture of 2% Isoflurane and O_2_ gas and depth of anesthetize was monitored by toe pinch throughout surgery. Prior to surgery, animals were weighed and given subcutaneous injections of the analgesic Meloxicam SR (4.0 mg/kg, Zoopharm) and saline (1 mL, Nurse Assist). Each fallopian tube was cauterized and bisected through lumbar inscitions on each side of the body using standard sterile practices. Animals were monitored once a day, for three days and staples were removed one week post-surgery. Females were given 2 weeks to recover before pairing with a male.

#### Pairing

For the RNAseq portion of the experiment, opposite-sex, non-sibling pairs ranged in age between P98 and P168 (average ∼P122) and same-sex, sibling pairs ranged in age between P60 to P163 (average ∼P111) at pairing. For the behavioral experiments, opposite-sex, non-sibling pairs ranged in age between P60 and P141 (average ∼P85) and same-sex, sibling pairs range in age between P62 and P79 (average ∼P76) at pairing. For RNAseq and behavior testing, at pairing, pairs were transferred into smaller static rodent cages (11.0 in. x 8.0 in. x 6.5 in.) with *ad libidum* water and rabbit chow (5326-3 by PMI Lab Diet), a cotton nestlet, and an igloo. Females were not induced. Partners cohabitated undisturbed, except for weekly cage changing performed by the experimenter, for 2 weeks prior to the baseline partner preference test (PPT). For same-sex pairings, sibling pairs from the same homecage were moved to smaller static rodent cages in the same way as the opposite-sex pairs. After the baseline PPT, behavior experiment animals were separated into individual cages for the remainder of the experimental timeline. In contrast, after the baseline PPT, RNAseq experimental animals were either returned to their partner (Remain Paired condition) or were separated into individual cages (Separated condition).

#### Separation for behavioral tests

Immediately following the baseline PPT, each pair was separated into fresh small static rodent cages (11.0 in. x 8.0 in. x 6.5 in.) with *ad libidum* water and rabbit chow (5326-3 by PMI Lab Diet), a cotton nestlet, and an igloo. Animals were placed approximately a foot away from each other to eliminate visual cues of their partner. Each animal in the pair remained in their individual cages undisturbed for either 48 hours (short-term separation) or 4 weeks (short-term separation) except for weekly cage changes done by the experimenter. At each separation time point, experimental animals performed a partner preference test (PPT) and resident intruder test (RIT) on successive days approximately 24 hours apart starting between 9:00 and 10:00 AM each day. Animals were returned back to their original home cages between testing days and transferred to fresh cages 24 hrs after RIT for each time point. Cage changes were done at identical intervals for each cohort prior to behavioral testing.

#### Partner Preference Test (PPT)

PPT was carried out as described by TH Ahern to assess selective partner affiliation prior to partner separation and were performed between the hours of 9:00 and 13:00 each testing day (1). Each 3 hour test was recorded by overhead cameras (Panasonic WVCP304) that film two boxes simultaneously. The test animal is placed in the middle chamber for 10 minutes with the cage dividers still in place. The test begins when the cage dividers are removed and the test animal is allowed free range of all three chambers for the duration of the test. The movement of the test animal was tracked post-hoc using Topscan High-Throughput software (v3.0, Cleversys Inc.) and frame by frame behavioral data was analyzed using a custom Python script (https://github.com/donaldsonlabdonald) to calculate the average distance between the test animal and tethered animals when in the same chamber, time spent huddling with each tethered animal, and total distance traveled. This data was then used to generate a number of behavior metrics including the partner preference score (Partner Huddle/Partner + Novel Huddle) (https://Fig.share.com/authors/zoe_donaldson_colorado_edu_Donaldson/6883910).

#### Resident Intruder Test (RIT)

Resident males were tested in their home cages (11.0 in. x 8.0 in. x 6.5 in.). Food hoppers and igloos were removed prior to the start of the test to allow video recording from the side using Sony Handycams (DCR-SX85). Intruder males were briefly anaesthetized with a mixture of ∼2% Isoflurane to shave their backs for identification during behavior scoring and allowed to fully recover in an empty cage. Intruders were then placed in the resident’s home cage and their behavior was filmed for 10 minutes. After the test, the intruders were returned to their home cages and the residents were returned their food hopper and igloos in their original home cage. Behavior scoring was done in BORIS (v7.10.2) by the same blind observer for all RIT tests. Behaviors scored included tumble fighting (aggression), chasing (aggression), defensive postures (aggression), anogenital sniffing (investigation), social investigation (investigation), huddling (affiliation), autogrooming (non-social), and digging (non-social). Definitions of scored behaviors can be found in **Table S1**. Behavior scores were then analyzed in R (v4.0.4).

#### Statistical analysis for behavioral experiments

All statistical analyses were performed in R (v4.0.4) using the base t.test package, base aov package, and the survival package (v3.2-7). Visualization was done using ggplot2 (v3.3.3), ggpubr (v0.4.0), and Adobe Illustrator (25.2.3). All statistical results can be found in **Table S1**. For PPT, we first determined if partner preference significantly changed from 50% at each time point (baseline, ST, or LT) and for each pairing type (OS or SS) by using an unpaired, one-sided Student’s t-test against a null value of 50. We also performed a 2-way repeated measures ANOVA with Tukey’s post-hoc test to determine main effects of, and interactions between, the pairing type and separation duration. Next, we calculated partner preference change scores for all conditions and compared the variance between pairing types using a F-test of variance. Finally, we determined changes to partner and novel huddling latency using the built in log-rank test of the survival package. For RIT, we determined any significant changes to the duration or bouts of all scored behaviors and behavior categories using a 2-way repeated measures ANOVA with Tukey’s post-hoc test. Additionally, to determine the latency to resident tumble fighting we used the built in logRank test of the survminer package (v0.4.8). For all behavior, we reported p-values that met a <0.10 threshold in order to highlight potential trends in behavior even if they did not meet the traditional p < 0.05 threshold.

#### Generation of vole optimized hSyn-DIO-eGFP-RPL10a construct for vTRAP

Prior to generating the vTRAP viral construct, the nucleotide and amino acid sequences for RPL10a in mice and prairie voles were compared. To assess nucleotide homology, the mouse RPL10a cDNA sequence was aligned to the vole RPL10a cDNA sequence using BLASTN (v2.8.033) with a percent identity of 93% and an E-value of 0.0 (**Fig. S3A**). The mouse RPL10a cDNA sequence was input into BLASTX (v2.8.033), and the resulting protein alignment against the prairie vole RPL10a amino acid sequence was 100% identical with an E-value of 3E-145 (**Fig. S3B**). Although the amino acid sequences are identical between mice and prairie voles, we generated the vTRAP construct using the prairie vole RPL10a cDNA sequence to account for possible species-specific differences in aminoacyl-tTRNA concentrations.

To generate the TRAP construct, the DNA fragment containing the vole RPL10a sequence was inserted into the pAAV-hSyn-DIO-eGFP (Addgene plasmid #50457) backbone. The RPL10a fragment was designed for cloning using the NEBuilder HiFi Assembly method and consisted of a 5’ homology arm sequence, a linker sequence coding for SGRTQISSSSFEF, the vole RPL10a sequence (GenBank XM_005360358.1), and a 3’ homology arm sequence (Supplementary Fig.. 1). pAAV-hSyn-DIO-eGFP was digested using restriction endonucleases Asc1 (NEB) and BsrG1-HF (NEB), and the linearized plasmid was purified using the Zymogen Gel DNA Recovery Kit. The cloned plasmid, hereafter referred to as pAAV-hSyn-DIO-eGFPpvRPL10a (vTRAP),was generated using the NEBuilder HiFi DNA Assembly Cloning kit and subsequently transfected into 5-alpha F’Iq Competen tE. Coli (NEB). Transfected bacteria were selected for using carbenicillin (100ug/ml) and harvested using the EZDNA Plasmid Mini Kit (Omega) or HiSpeed Plasmid Maxi Kit (Qiagen). The sequence of vTRAP was verified using Sanger sequencing (QuintaraBio) and restriction digest before being sent for viral packaging in an AAV1 serotype (Stanford Neuroscience Gene Vector and Virus Core).

#### Viral injections

For stereotaxic surgery, adult prairie voles were anesthetized using isoflurane (3% induction, 1-2.5% maintenance) and depth of anesthesia was monitored by breathing and toe pinch reflex. After induction, animals were placed on a heating pad and given an analgesic (2 mg/kg Meloxicam SR, subcutaneous, Zoopharm) and sterile saline (1 mL, subcutaneous). Standard sterile surgical procedures were followed to expose and drill through the skull. Nucleus accumbens injection coordinates were adapted from the Allen Mouse Brain Atlas and validated prior to the experiment (AP: -1.7, ML: ±1.0, DV: -4.7, -4.6, -4.5, -4.4, -4.3). For each DV coordinate, 100 nL of virus was dispensed at 0.1 nL/sec for a total of 500 nL using a Nanoject II (Drummond Scientific). After the last injection, the needle remained in place for 10 minutes to allow for diffusion. The glass needle was then removed and the skin was sutured closed (vicryl sutures, eSutures), coasted with lidocaine and antibacterial ointment, and animals recovered in a cage on a heating pad with continued monitoring. The animal’s health was monitored for 3 days following surgery and they were allowed to recover in their homecage for 2 weeks prior to the experiment start.

#### Verification of TRAP Expression

Cre-dependent expression of the vole optimized eGFP-RPL10a custom virus was validated in adult male prairie voles (n = 1) using a within animal design. The left NAc was injected with a viral cocktail of AAV1-DIO-eGFP-RPL10a (2 uL; 5.55E11 vg/mL) and AAV1-hSyn-iCre (2 uL; 3.9E12 vg/mL) while the right NAc only received the AAV1-DIO-eGFP-RPL10a virus (600 nL; 2.22E+12 vg/mL). Following 2 weeks of recovery to allow for robust viral expression the animal was perfused. The animal received a lethal injection of 0.2 mL 1:2 ketamine/xylazine and was perfused with 1X PBS followed by 4% paraformaldehyde (in 1X PBS, Electron Microscopy Sciences) and kept in 4% PFA overnight. The brain was then placed in 5 mL of 30% sucrose for 3 days prior to sectioning on a microtome (Leica JungSM2000R, 50 microns/slice). Sections were wet mounted onto Superfrost Plus glass slides (Thermo Fisher Scientific), cover slipped (ProLong Gold; Invitrogen), and allowed to dry overnight prior to imaging. Prepared slides were then sealed with nailpolish (Electron Microscopy Sciences) and imaged using a Yokogawa CV1000 Confocal Scanner System (University of Colorado Light Microscopy Core Facility). Stitched images were produced using ImageJ (v1.51).

#### Separation for RNAseq experiments

Immediately following the baseline PPT, each pair was moved to fresh small static rodent cages (11.0 in. x 8.0 in. x 6.5 in.) with *ad libidum* water and rabbit chow (5326-3 by PMI Lab Diet), a cotton nestlet, and an igloo. Remain Paired animals were kept with their partner while Separated animals were separated from their partner and moved to a fresh cage while their partner was immediately SAC’d to remove confounds of visual or olfactory cues. All animals remained in their individual cages undisturbed for either 48 hours (acute separation) or 4 weeks (chronic separation) except for weekly cage changes done by the experimenter. After the appropriate separation time point, experimental animals were removed directly from their home cage for rapid decapitation and tissue harvesting.

#### RNA preparation and sequencing

Tissue lysis and RNA isolation was adapted from Heiman and performed under RNAse free conditions (2). On each tissue collection day, 2 animals from each experimental cohort were processed, with the exception of same-sex 4 week paired animals who were all collected on the same day. Animals were removed from home cages, rapid decapped, and both hemispheres of the Nucleus Accumbens were hand dissected using sterile, RNAse free razor blades and pooled. During dissection, tissue was kept wet with dissection buffer (HEPES KOH, pH 7.3 20 mM, 1X HBSS, Glucose 35 mM, NaHCO3 4 mM, CHX 10 uL of 1000X stock, DEPC-treated water up to 10 mL). Tissue was added to 1 mL of ice-cold lysis buffer (DEPC-treated water, HEPES 20 mM, KCl 150mM, MgCl2 (10 mM), DTT 5 uL of 1M stock, CHX 10 uL of 1000X stock, 1 protease inhibitor, RNasin 400 Units, and Superasin 800 Units), immediately homogenized using a Scilogex homogenizer, and kept on wet ice while the remaining animals for the day were processed. The homogenizer tip was thoroughly cleaned in between animals with DEPC-treated water and 70% ethanol to eliminate cross contamination between samples. Homogenized samples were centrifuged at 4°C, for 10 minutes at ∼3000Xg and supernatent was transferred to new, pre-chilled tubes. 10% NP-40 (AG Scientific: #P-1505) and 300 mM DHPC (Avanti Polar Lipids, Inc.: 850306P-200 mg) were added to the supernatent and incubated on ice (4°C) for 5 minutes. Samples were then centrifuged at 4°C for 10 minutes at ∼16000Xg and supernatent was transferred to new, pre-chilled tubes. RNA was then extracted and cleaned from the lysate supernatent using the Norgen Total RNA micro kit (cat. #) according to the manufacturer’s instructions. RNA integrity was measured using an Agilent Tapestation prior to library preparation. RINs for all samples were >7.3 (average 8.8).

For TRAP samples, dissections and Input fraction RNA were prepared as above. To perform immunoprecipitation pulldowns, affinity matrices were prepared using 300 uL of 10 mg/mL Streptavidin MyOne T1 Dynabeads (Invitrogen), 150 uL of 1ug/uL Biotinylated Protein L (Fisher Scientific), and 50 ug each of eGFP antibodies Htz-19C8 and Htz-19F7 (Memorial-Sloan Kettering Monoclonal Antibody Facility) 2 hours prior to adding Input RNA sample. A small sample of Input RNA was saved for next day clean up (∼250 uL/sample) and frozen at -80C overnight. The remaining Input RNA (∼1 mL) was added to the GFP-conjugated beads and rotated overnight (16-18 hours) at 4 °C. The next day, the flowthrough was saved and stored at - 80C and bound mRNA was released from the beads using a series of high salt buffer washes (4 washes; 20mM HEPES, 150mM KCl, 10mM MgCl2, 10% NP-40, RNase-free water, protease inhibitor tablets, 0.5mM DTT, 100ug/ul cycloheximide). The beads were then incubated with the Norgen Total RNA micro kit SKP buffer (300 uL) for 10 minutes at room temperature. The Pulldown supernatant was then saved and processed as above using the kit manufacturer’s instructions.

Library preparation for sequencing was done using the KAPA mRNA HyperPrep kit with polyA enrichment according to the manufacturer’s instructions for a 100-200 bp insert size and using 1.5 uM TruSeq3-SE adapters. 71 samples were pooled and 3 runs of NextSeq V2 high output 75 cycle (1×75) sequencing produced a total read depth of ∼20M reads/sample. The Illumina sequencing was performed at the BioFrontiers Institute Next-Gen Sequencing Core Facility of University of Colorado Boulder. RNA quantity, RIN, and RNA amount used for library preparation can be found in **Table S1**.

#### Sequence mapping and counting

For quality control, FastQC (0.11.8) and Trimmomatic-Src (v0.38) were used to analyze read quality and to trim the TruSeq3-SE adapters from all samples (3). Additionally, each sequencing run was correlated to each other to ensure that there was not a significant difference in run composition. Reads were aligned to the prairie vole genome (Microtus_ochrogaster.MicOch1.0.dna.toplevel, released Feb 2017; accession GCA_000317375) using HiSat2 (2.1.0) with default options and individual runs for each animal were merged into a single sample using SAMtools (1.9) (4, 5). Reads were then counted using HTseq (0.11.2) with default options using the annotated prairie vole genome (Microtus_ochrogaster.MicOch1.0.95.gtf, released Feb 2017; accession GCA_000317375) (6). Genes with less than 10 read counts were filtered from downstream analysis and, where necessary, were normalized using transcripts per million (TPM). Where possible, all Ensemble IDs were converted to prairie vole gene IDs using BioMart. If no gene ID was found, the original Ensemble ID was retained. Only genes with greater than 10 counts were included in subsequent analyses (total genes = 21152, genes >10 counts = 12128).

#### Data analysis and visualization

All sequencing analysis was performed using R (v4.0.4) (7). The packages used and their specific parameters are described in the relevant sections below. Plots were made using base plot, ggplot2 (v3.3.3), EnhancedVolcano (1.8.0), RRHO2 (v1.0), clusterProfiler (v3.18.1), and Adobe Illustrator (25.2.3) (8–11).

#### DESeq2

Differential gene expression was analyzed with the R package DESeq2 (v1.30.1) following the standard workflow (12). To determine differential gene expression between cohorts a DESeq dataset of all animals was made with the design ∼ParentCode + AgeAtPairing + Cohort to correct for the effect of parentage and age at initial pairing. Other batch effects (processing day, processing order, RIN, etc.) did not have sufficient influence on the model to include. Each result was a pair-wise contrast between the appropriate cohorts with the opposite-sex condition as the experimental and the same-sex condition as the reference. Differential gene expression was defined as >0.30 and <-0.30 log2FoldChange and p-value <0.05. This Log2FoldChange threshold was used for two reasons. 1) A threshold of +/- 0.20 can be successfully validated by qPCR (13) but 2) we chose a slightly higher threshold as it allows for smaller gene lists that appear more biologically relevant by Gene Ontology analysis. Differential gene expression was visualized using EnhancedVolcano. To determine if there was statistically significant overlap between differentially expressed gene lists compared to all genes (n = 12,128) we performed a Fisher’s exact test using the SuperExactTest (v1.0.7) package (14). Additionally, we used Morpheus (https://software.broadinstitute.org/morpheus) heatmaps to compare the expression profiles of each DESeq comparison using scaled log2FoldChange values. For each heatmap, a reference dataset ordered the remaining datasets such that each column is the same gene and the color represents the scaled log_2_FoldChange in each DESeq comparison. In *Fig. 2F* the short-term pair bond was set as the reference and in *Fig. 3F, J*, and *L* the combined pair bond was set as the reference. Gene ontology and pathway analysis of relevant gene lists were performed using the R package enrichGO with *Mus musculus* ontology terms (11). Representative terms with an adjusted p-value of <0.05 were hand selected. Results were visualized using base R plots modified in Illustrator. All DESeq result data frames can be found in **Table S2**.

#### Rank-Rank Hypergeometric Overlap (RRHO)

RRHO is a threshold free approach used to identify genome-wide expression patterns between two ranked gene lists. RRHO analysis was performed using the RRHO2 (v1.0) package with parameters of stepsize = 100 and boundary = 0.02. First, transcripts from each DESeq comparison are ranked by their p-value and effect size direction:

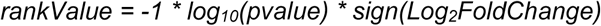

This results in a ranked list where the most significantly upregulated genes are at the top of the list and the most significantly downregulated genes are at the bottom of the list with each gene appearing only once. Next, two ranked gene lists are compared to each other using a hypergeometric distribution to determine the significance of the number of overlapping genes within defined sample populations. Sample populations are independently defined for each list. Each population consists of all of the genes that are ranked higher than a specified threshold. Thresholds successively move down a list at a user specified step size. For computation convenience, here we define our step size as 100 genes. However, the smallest possible step size is 1 gene and the largest possible step size is the number of genes found in the smallest list. The hypergeometric p-values resulting from comparing the List 1 genes (all genes above threshold x) to List 2 (all genes above threshold y) then populate a square matrix. As a result, the furthest bottom-left position of the matrix is the p-value of the overlap between genes ranked above the 1st threshold in each list and the furthest top-right position is the overlap between genes ranked above the last threshold in each list. As another example, the furthest bottom-right position is the p-value comparing all genes above the last threshold of List 1 to all genes above the 1st threshold of List 2. The final matrix of hypergeometric p-values is visualized as a heatmap where List 1 is along the x-axis and List 2 is along the y-axis and each point represents a p-value. The axises are arranged such that genes that are upregulated in both lists are in the bottom-left quadrant (quadrant UU) and genes that are downregulated in both lists are in the top-right quadrant (quadrant DD). Genes with opposite regulation in each condition, up in one list but down in the other, are along the opposite diagonal (quadrants UD and DU) (Fig.ure 2E). To visualize RRHO our heatmaps with a consistent p-value scale we set the scale maximum = 300 and the minimum = 1.

The genes in each quadrant were extracted to form UU, DD, UD, and DU gene lists for further analysis. Using these gene lists, the significant overlap between quadrants were determined using the SuperExactTest package. Gene ontology and pathway analysis of these gene lists were performed using the R package enrichGO with *Mus musculus* ontology terms. Representative terms with an adjusted p-value of <0.05 were hand selected.

Additional information for RRHOs are in **Fig. S2**. Unscaled p-value RRHOs can be found in **Fig. S2A-C**. Since the p-value of the hypergeometirc distribution is sensitive to the number of genes, RRHO also plots a log-Odds ratio for all comparisons. Log-Odds ratio plots can be found in **Fig. S2D-F** where the maximum scale was changed from 8 to 6 on all plots to better visualize values. Finally, to ensure that the signal we see in the RRHOs is not due to chance, we shuffled the ranked order of List 1 in each RRHO (**Fig. S2G-I**). The resulting lack of signal throughout the RRHOs support that the signal we see in our experimental RRHOs is not due to chance.

#### Pathway and Upstream Regulator Analysis with Ingenuity Pathway Analysis (IPA)

Ingenuity pathway analysis (Qiagen, Germantown, MD), was used to determine potential biological pathways and upstream regulators of DEGs and co-regulated genes identified using RRHO. Core expression analysis was applied for each gene list followed by comparison analysis for related lists to help identify common pathways and regulators across groups. For both, outcomes were filtered to include only those that reached an activation z-score of +/-2 and a corrected p-value of <0.05 (corrected by Benjamini Hochberg). All predicted biological pathways and upstream regulators are presented in **Supplemental Table 3**.

#### Weighted Correlation Network Analysis (WGCNA)

A weighted gene co-expression network analysis was conducted using the R package WGCNA (v1.70-3) (15). WGCNA analysis was done using the Input and Pulldown sequencing data from only animals in the vTRAP cohorts (opposite-sex remain paired 4 weeks, opposite-sex separated 4 weeks, and same-sex separated 4 weeks). A signed co-expression network was constructed using a power of 9 and minimum module size of 30 genes according to the author’s standard module-trait relationship workflow. No samples were designated as outliers using complete clustering. Gene modules were then correlated to either the trait Input or Pulldown to determine the gene clusters associated with each sample type. To determine which correlation direction (positive or negative) is associated with which sample type, the significantly positively or negatively correlated modules were grouped and analyzed by Enrichr for cell types (16–18). The top term for the positively correlated modules was nucleus accumbens while the top term for the negatively correlated modules was spinal cord. Therefore, the positively correlated modules represent the Pulldown samples, and by extension, neuronally enriched genes.

#### Neuronal enriched genes throughout separation

To robustly define neuronally enriched genes we found the intersection of the vTRAP Input vs Pulldown enriched DEGs (**Fig. 4B**) and the genes from the significantly, positively correlated WGCNA modules (**Fig. 4D**). The resulting intersection—2,365 genes— is referred to as “neuronal enriched genes” and includes almost all of the vTRAP enriched DEGs. To further validate the neuronal enriched genes we analyzed this gene list using Enrichr for cell types. The top two terms are nucleus accumbens and dorsal striatum confirming the specificity of this gene list.

We next wanted to follow the expression of these neuronal enriched genes throughout separation. First, since we only had vTRAP samples from specific cohorts in our experiment, we ensured that neuronal gene expression can be inferred from bulk sequencing of the same genes. To validate this, we filtered the Input and Pulldown samples of the opposite-sex 4 week separated cohort and compared the expression of the neuronal enriched genes using Morpheus (Fig. S5C). The strong similarity in expression of the neuronal enriched genes using two different collection methods validated that bulk sequencing expression can be used to infer the neuron specific expression of those same genes. Second, to compare the expression patterns of the neuronal enriched genes to the bulk sequencing data we filtered the bulk data for only the neuronal genes. We then clustered the unfiltered bulk sequencing data, from *3F*, and the neuronal enriched genes only bulk sequencing data using one minus Pearson correlation with complete linkage. For each neuronal cluster we then found the corresponding cluster in the unfiltered expression data with the correspondence links visualized by a Sankey plot using the R package networkD3 (v0.4) (http://christophergandrud.github.io/networkD3/). Third, we identified three clusters in the neuronal enriched data that had differential expression based on separation duration. We analyzed each gene list using Gene Ontology as previously described and highlighted specific genes associated with top terms.

**Fig. S1.**
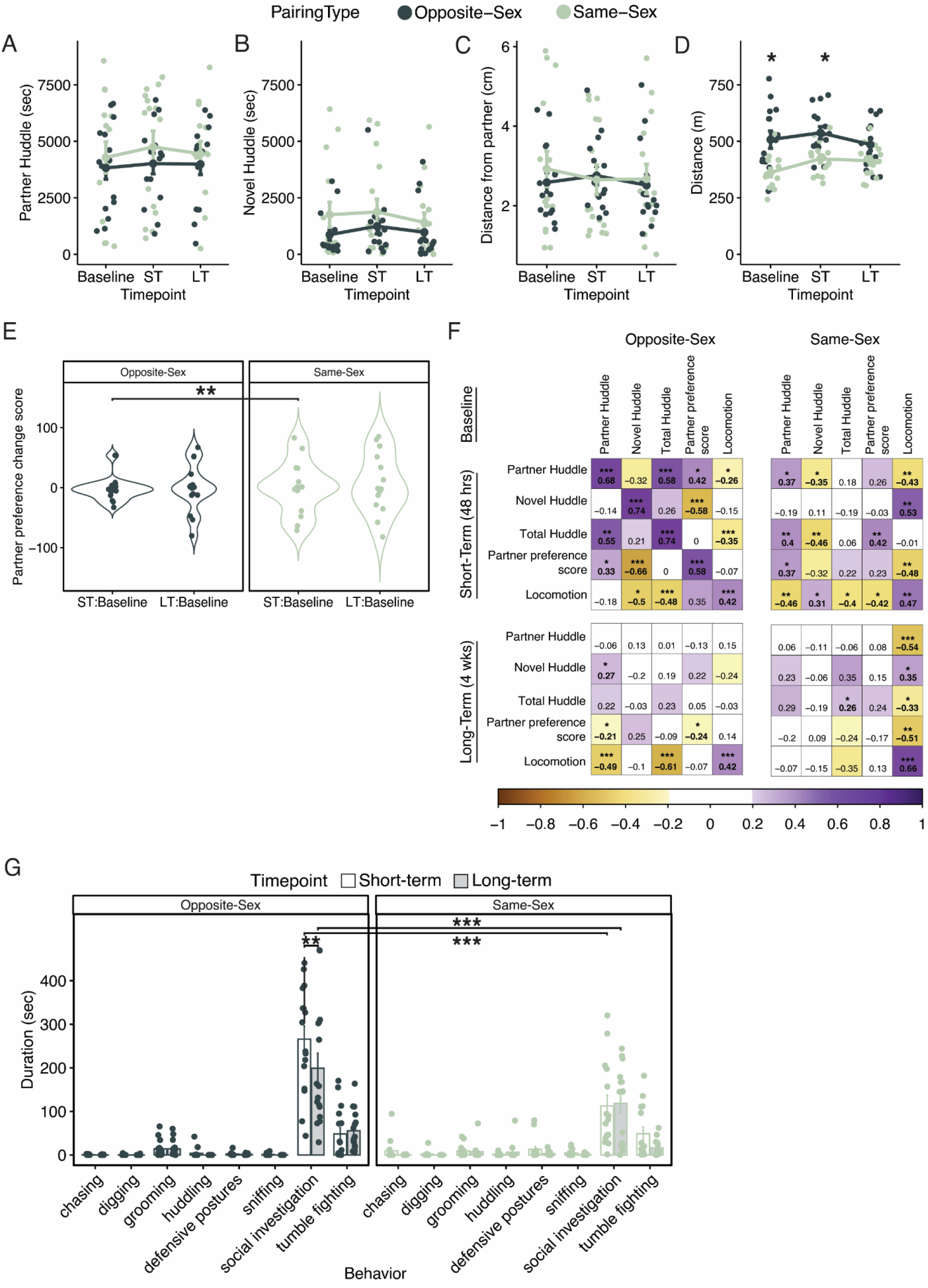
Opposite-sex- and same-sex-paired animals display subtle behavioral differences in the partner preference test and resident intruder test behaviors following separation. **(A-C)** For opposite-sex (OS; dark green) and same-sex (SS; light green) animals at baseline, short-term separation (ST) and long-term separation (LT), there is no significant difference in total partner huddle duration (sec) (A), novel huddle duration (sec) (B), or distance (cm) of the experimental animals from their partner (C). **(D)** OS-paired animals locomoted significantly more in the partner preference test than SS-paired animals during the baseline and short-term test but not during the long-term test (2-way RM-ANOVA with Tukey’s post-hocs: Baseline p = 0.0017, short-term p = 0.024, long-term p = 0.36). **(E)** Violin plots of partner preference change scores ST:Base (short-term minus baseline) and LT:Base (long-term minus baseline). Opposite-sex paired males had significantly less variance in their partner preference change scores between baseline and short-term separation than same-sex paired males (F-test of equality of variances: F_14_ = 0.2045, p = 0.0054) though the effect of pairing type is no longer significant after long-term separation (F_14_ = 0.673, p = 0.469). **(F)** Pearson’s correlation matrices of partner preference test behaviors across time points (Top: baseline, Middle: short term, Bottom: long term) in both pairing types (opposite-sex; left, same-sex;right). Number and color indicate Person’s R-value. Locomotion remains similarly correlated across all comparisons indicating that the changes in behavioral consistency between time point and pairing type is restricted to social behaviors. ***p < 0.001, **p < 0.01, *p < 0.05. **(G)** Bar graph of the duration (sec) of each behavior scored during the resident intruder at short-term (white) and long-term (grey) separation time points. There is a significant difference in social investigation between pairing types at both time points and in OS-paired animals between short- and long-term separation (3-way RM-ANOVA with Tukey’s post-hoc: ST opposite-sex vs same-sex p = 9.3 × 10^−12^; long-term opposite-vs same-sex p = 3.2 × 10^−4^; opposite-sex short-vs long-term p = 0.011).

**Fig. S2.**
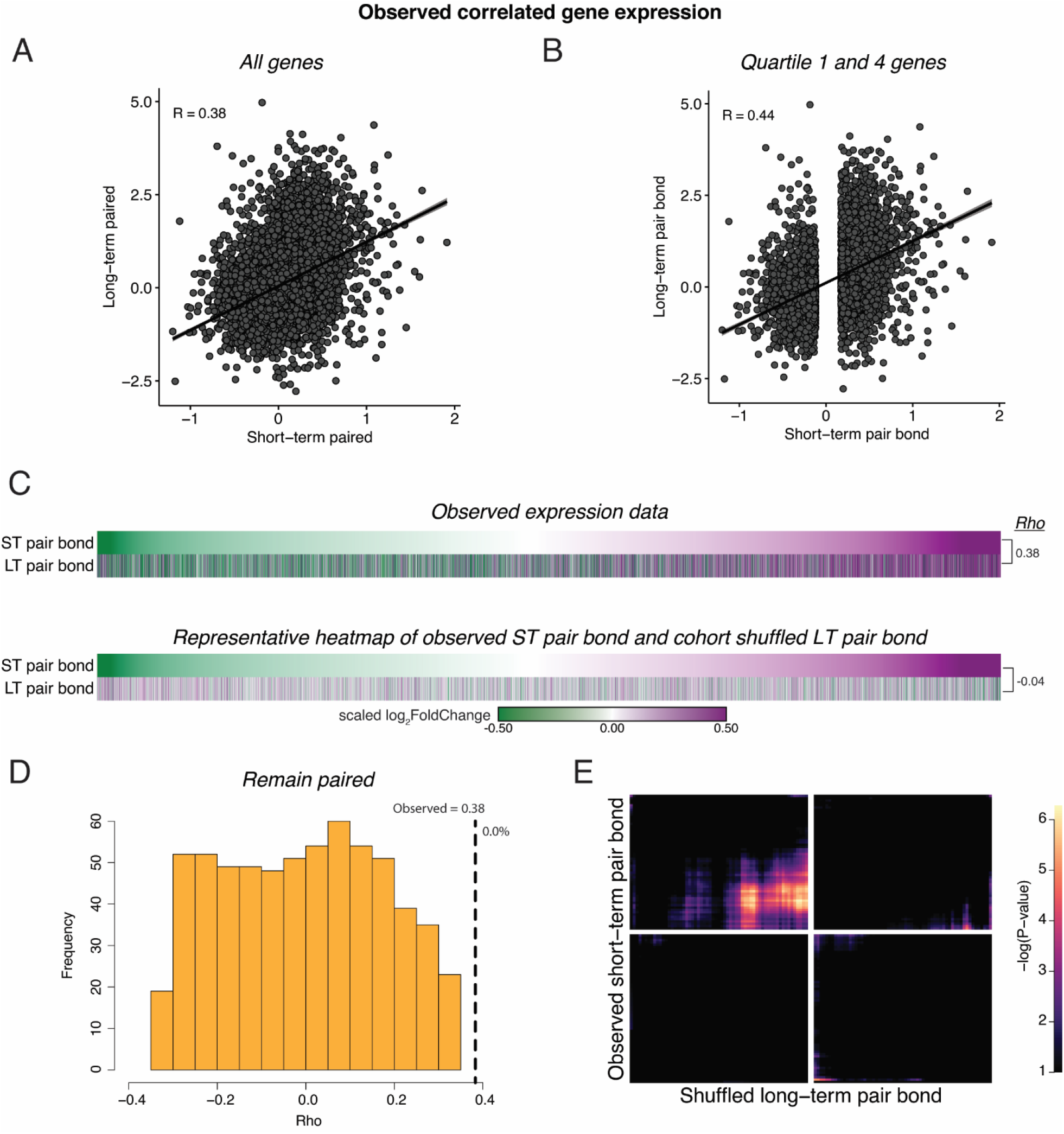
Cohort shuffled controls for remain paired cohorts. **(A)** Spearman’s Ranked Correlation of gene expression (log_2_FoldChange) at the short-term pair bond and the long-term pair bond. Black line is the regression line with shading indicating a 95% confidence interval. **(B)** To ensure that our correlations are not driven by genes with consistent but low differential expression values, we only retained the first and fourth quartile of genes in the short-term pair bond and found a stronger Spearman’s Ranked Correlation between pair bond time points. **(C)** To test if the correlation in gene expression between remain paired timepoints was likely to be due to chance, we shuffled the Cohort identity for each animal and performed differential expression analysis. Heatmaps show every gene from short-term and long-term pairing ordered from the smallest to largest log_2_FoldChange after short-term pairing in the observed comparison (top) and a representative Cohort shuffle (bottom). Scale represents the scaled log_2_FoldChange. **(D)** Histogram of Rho values for 1000 iterations of (C, shuffled data). Only correlations that did not have a –log_10_(adjusted pvalue) of infinite were retained (n = 636). Vertical dashed line indicates the observed Rho value with the percent of shuffled iteration in which Rho was greater than the observed value to the right (0.0%). **(E)** Unscaled RRHO plot of the representative Cohort shuffle used in (C) indicates elimination of the concordant signal seen with the observed data.

**Fig. S3.**
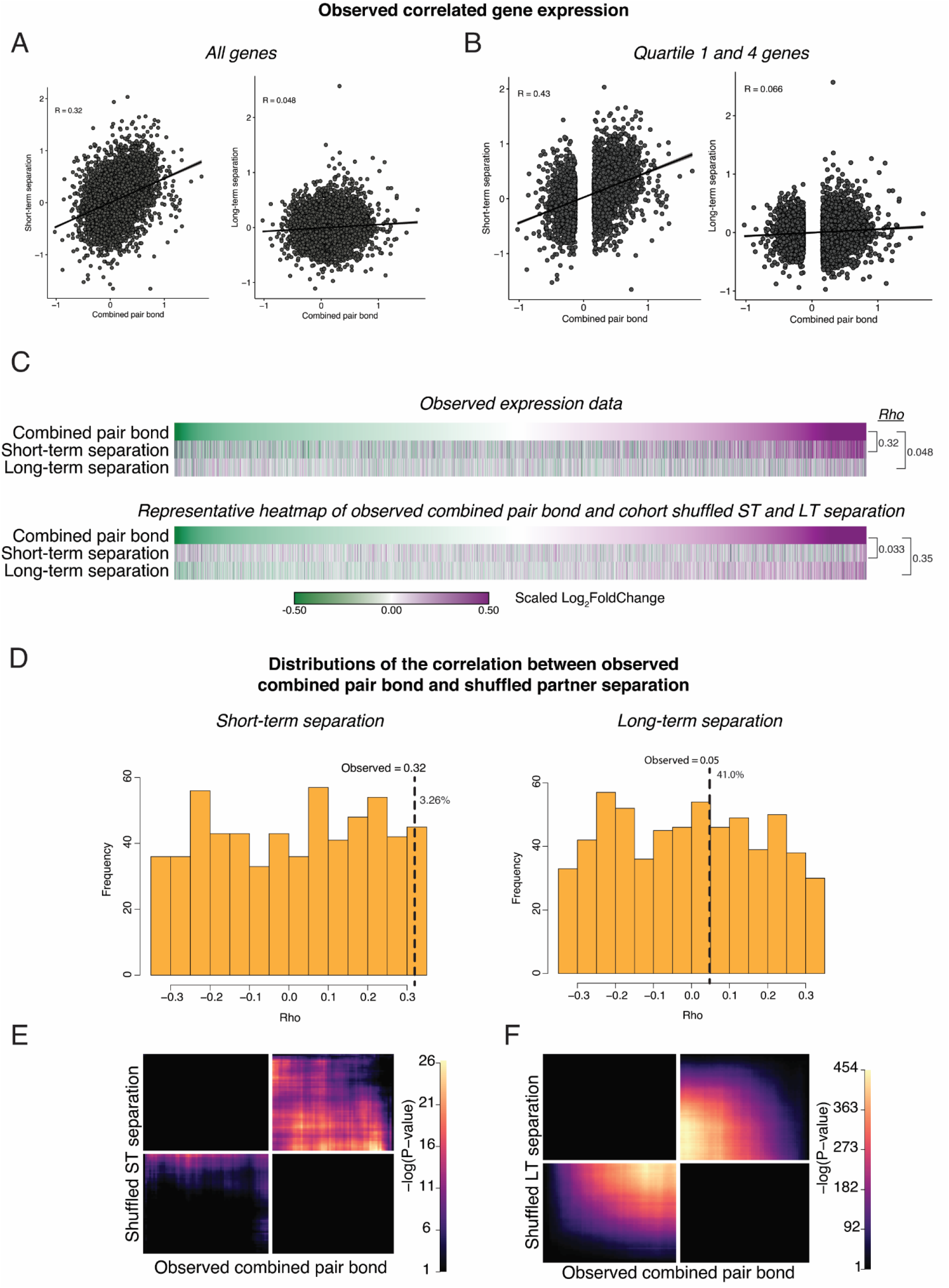
Cohort shuffled controls for the combined pair bond and separation time points. **(A)** Spearman’s Ranked Correlation of gene expression (log_2_FoldChange) for the combined pair bond vs short-term separation (left) or the combined pair bond vs long-term separation (right). Black line is the regression line with shading indicating a 95% confidence interval. **(B)** To ensure that our correlations are not driven by consistent but low differential expression values we only retained the first and fourth quartile of genes in the combined pair bond. We then performed a Spearman’s Ranked Correlation between the combined pair bond and each separation time point and found a greater Rho value for each time point. **(C)** To test if the correlation in gene expression between timepoints could occur by chance we shuffled the Cohort identity for each animal and performed differential expression analysis. Heatmaps show every gene from the combined pair bond and separation time points ordered from the smallest to largest log_2_FoldChange in the combined pair bond for the observed comparison (top) and a representative Cohort shuffle (bottom) with Rho values for each. Scale represents the scaled log_2_FoldChange. **(D)** Histogram of Rho values for a 1000 shuffles of the Cohort identity as in C, shuffled. Rho values for the correlation between the combined pair bond vs short-term separation (left) or the combined pair bond vs long-term separation (right) are plotted for each iteration. Only correlations that did not have a –log_10_(adjusted pvalue) of infinite were retained (short-term separation n = 613; long-term separation n = 617). Vertical dashed line indicates the observed Rho value with the percent of shuffled Rho’s that were greater than the observed value to the right (pair bond:short term 3.26%; pair bond:long term 41%). **(E, F)** Unscaled RRHO plot of the representative Cohort shuffle used in (C). The combined pair bond vs short-term separation (E) indicates a similar concordant pattern as the observed data though to a much lower degree (note scale maximum). The combined pair bond vs long-term separation (F) shows a highly significant concordant pattern that does not occur in the observed data.

**Fig. S4.**
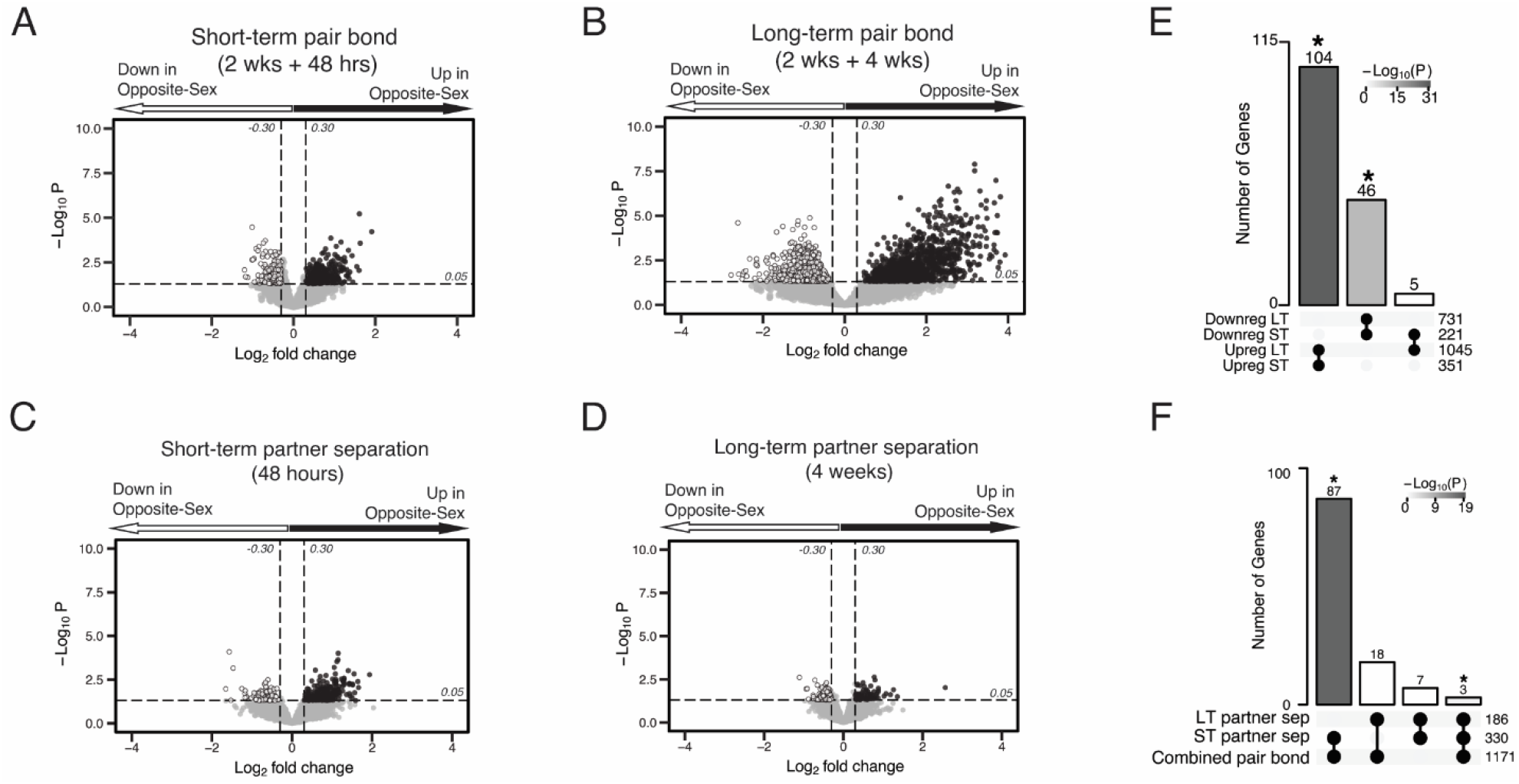
Volcano plots of differential expression with Upset plots for overlap of differential expressed genes. **(A-D)** Volcano plots with associated Upset plots of the differentially expressed genes of the short-term pair bond (A), long-term pair bond (B), short-term partner separation (C), and long-term partner separation (D). Genes that are upregulated in opposite-sex paired males compared to same-sex paired males are to the right (black dots) and genes that are downregulated are to the left (white dots). Dashed lines indicate thresholds of +/- 0.30 log_2_FoldChange (vertical) or 0.05 p-value (horizontal) used to determine differential expression. **(E)** Upset plot showing overlap of up- and downregulated DEGs from each timepoint. Intersections with no overlapping genes, such as downregulated long-term:upregulated short-term, are not displayed on the graph. Asterisks and shading indicate statistically significant intersections (Fisher’s Exact Test: Upreg ST:LT χ^2^= 30.24, p = 5.36 × 10^−31^; Downreg ST:LT χ^2^= 13.32, p = 7.05 × 10^−14^). **(F)** Upset plot showing the combined up- and downregulated separation DEGs compared to the combined pair bond DEGs. There are more shared transcripts between the pair bond and short-term separated animals than the pair bond and long-term separated animals. Asterisks and shading indicate statistically significant intersections (Fisher’s exact test: pair-bond:short-term sep χ^2^= 31.86, p = 6.78 × 10-19; pair-bond:long-term sep χ^2^= 17.96, p = 0.53).

**Fig. S5.**
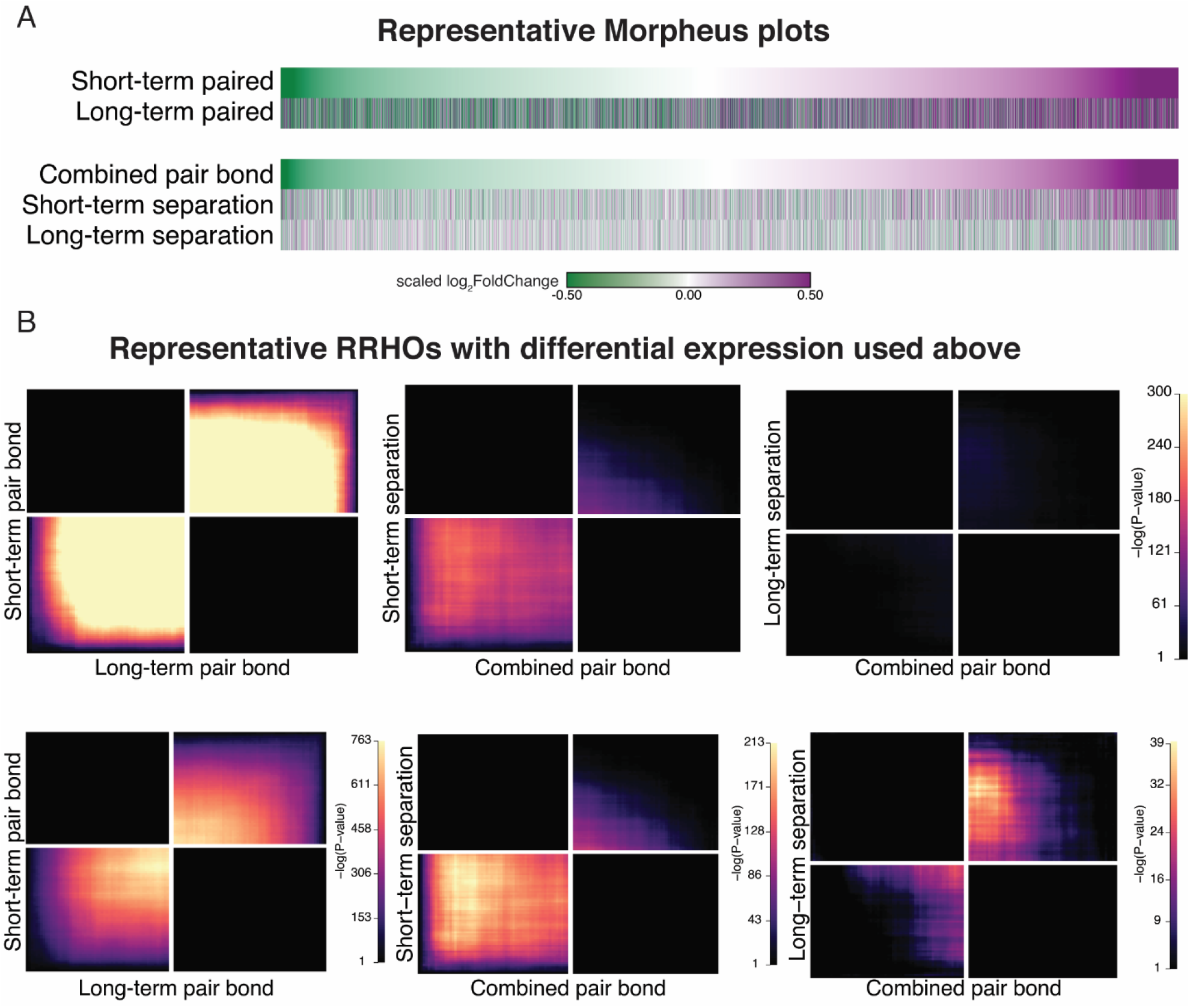
Randomly selected subsampled data shows the same differential expression and RRHO patterns as the full dataset. **(A)** We selected a random subset of the same number of animals for each opposite-sex (n = 6) and same-sex paired (n = 5) cohort, and performed differential expression analysis between pairing time points (top) and the combined pair bond vs separation time points (bottom). The heatmap shows genes ordered based on log2FoldChange in the short term Pair bond (tope) or combined pair bond (bottom) with color indicated expression difference between opposite and same-sex paired males. Scale represents the scaled log_2_FoldChange. **(B)** RRHO heatmaps of the representative subset used in (A) that use the same scale as the observed data (top) or are unscaled to visually optimize significant signal (bottom).

**Fig. S6.**
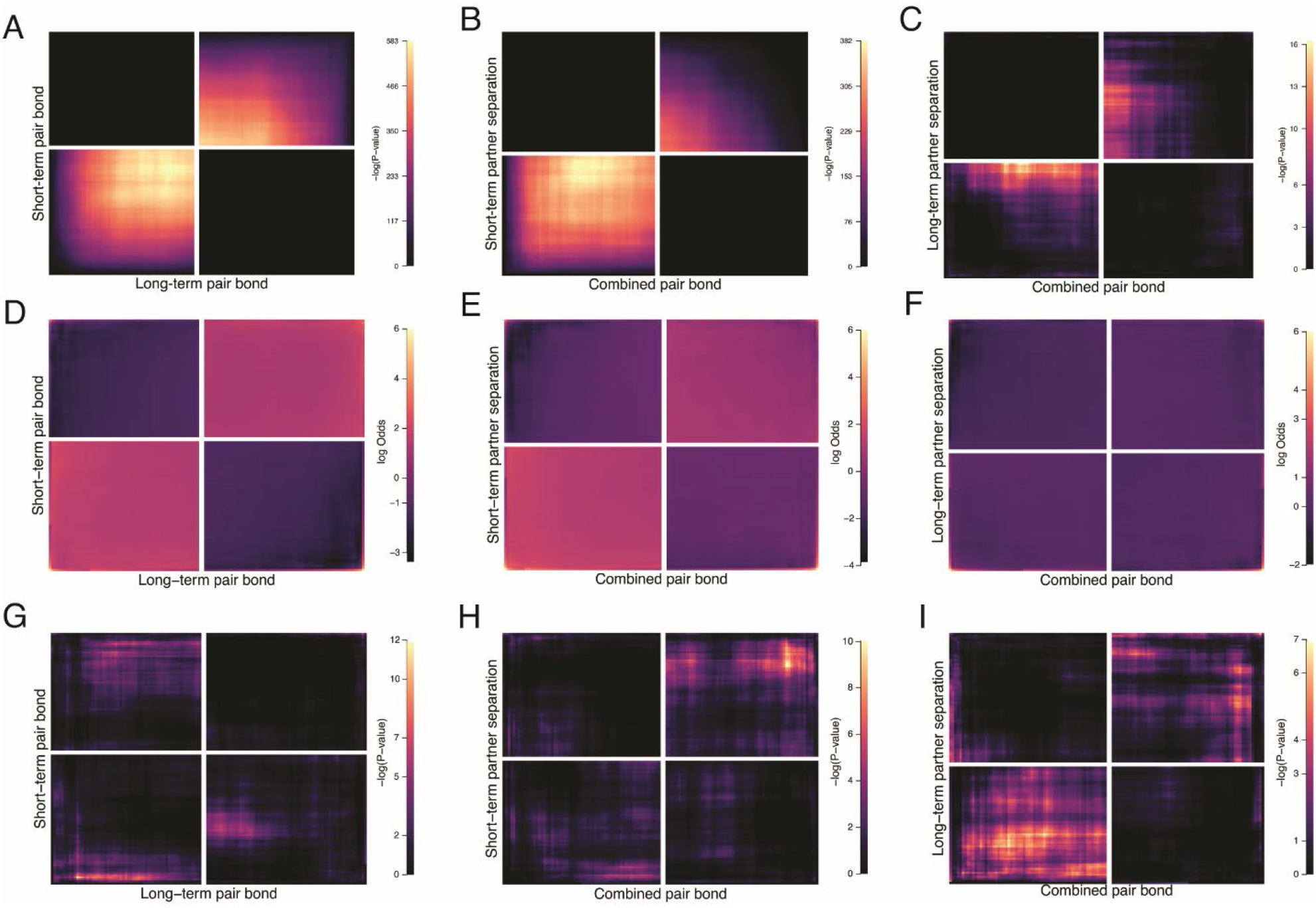
Control analyses for RRHO. Further examination of RRHO results where A, D, G correspond to *Figure 2F* (short-term vs long-term remain paired). B, E, F correspond to *Fig 3E* left (pair bond vs short-term separation). C, F, I correspond to *Fig 3E* right (pair bond vs long-term separation). **(A, B, C)** RRHOs showing - log_10_(p-value) scales optimized for visualization of significant signal. Note the greatly reduced scale (max 16) in *C* compared with other analyses. **(D, E, F)** Log-Odds ratio plots for RRHOs of all analyses scaled to a max value of 6. **(G-I)** Control analyses where the ranked gene lists were randomly shuffled while all other RRHO parameters remain the same.

**Fig. S7.**
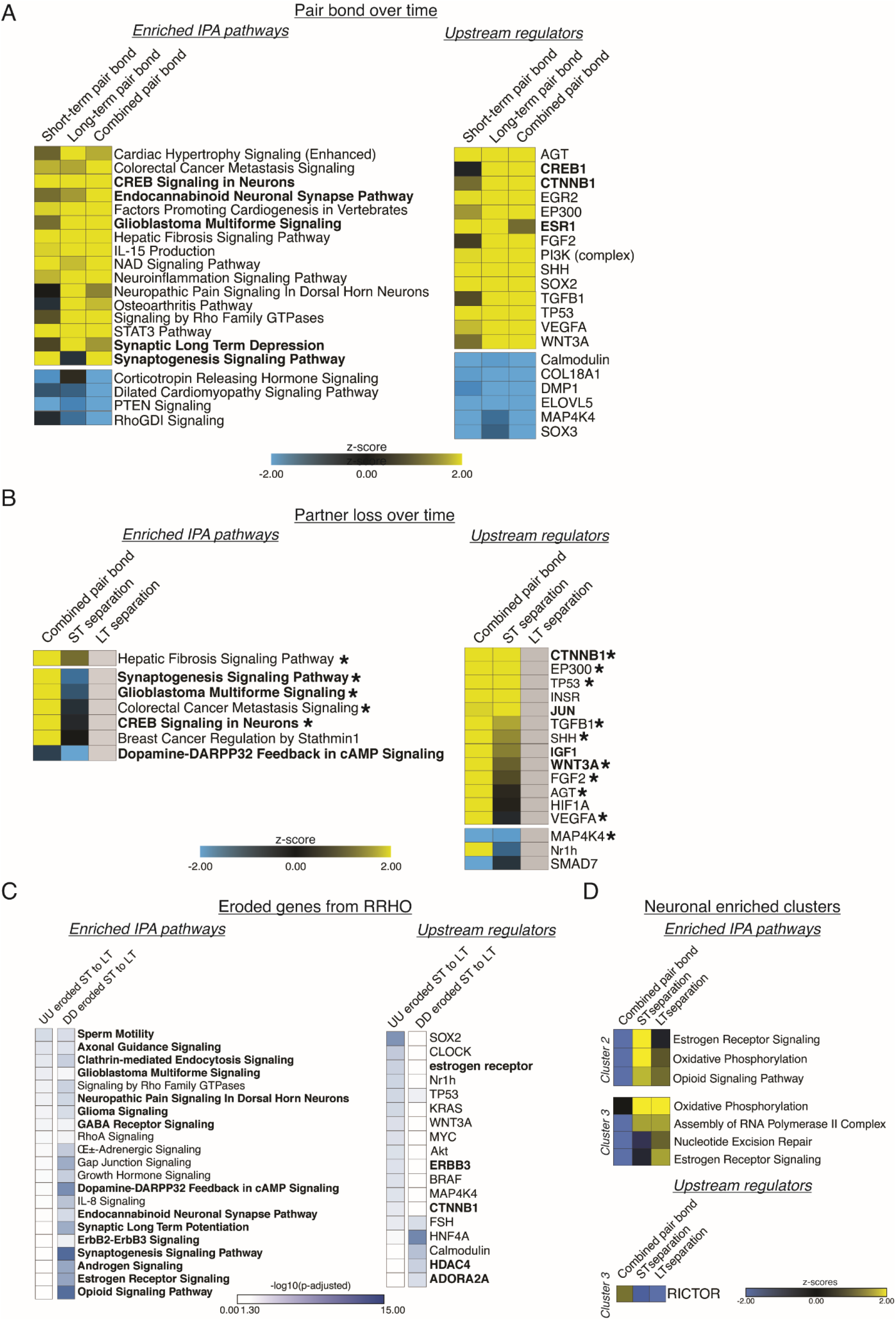
Ingenuity Pathway Analysis (IPA) corresponding to GO term analyses. IPA analyses indicating enriched pathways and upstream regulators for applicable comparisons. Representative pathways and regulators were chosen based on significance, activation pattern, and relationship to GO terms. Full lists can be found in Supplemental Table 3. Yellow to blue scales indicate the z-scores of enrichment where yellow is activated and blue is repressed. White to blue scale represents the –log_10_(p-adjusted) where any non-significant terms are white. **(A)** IPA analysis for the remain-paired cohorts in *Fig 2*. The strong similarity of the activation/repression of pathways and regulators from short-term to long-term pairing are well represented by the combined pair bond transcriptional signature. Pathways or regulators of interest are bolded. **(B)** IPA analysis of the combined pair bond, short-term separation, and long-term separation in *Fig 3*. Grey boxes indicate that the term is not found in the results for that time point. Terms and regulators of interest are bolded while asterisks denote terms or regulators found in A. **(C)** IPA analysis of the eroded ST to LT genes for the UU and DD quadrants of *Fig 3*. Results are denoted as adjusted p-values instead of z-scores because there is no log_2_FoldChange value associated with the RRHO heatmaps. Pathways or regulators of interest are bolded. **(D)** IPA analysis of the neuronal enriched clusters of interest from *Fig 4*. Cluster #1 contained few genes and no significant pathways or regulators. Cluster #2 did not have any significant regulators.

**Fig. S8.**
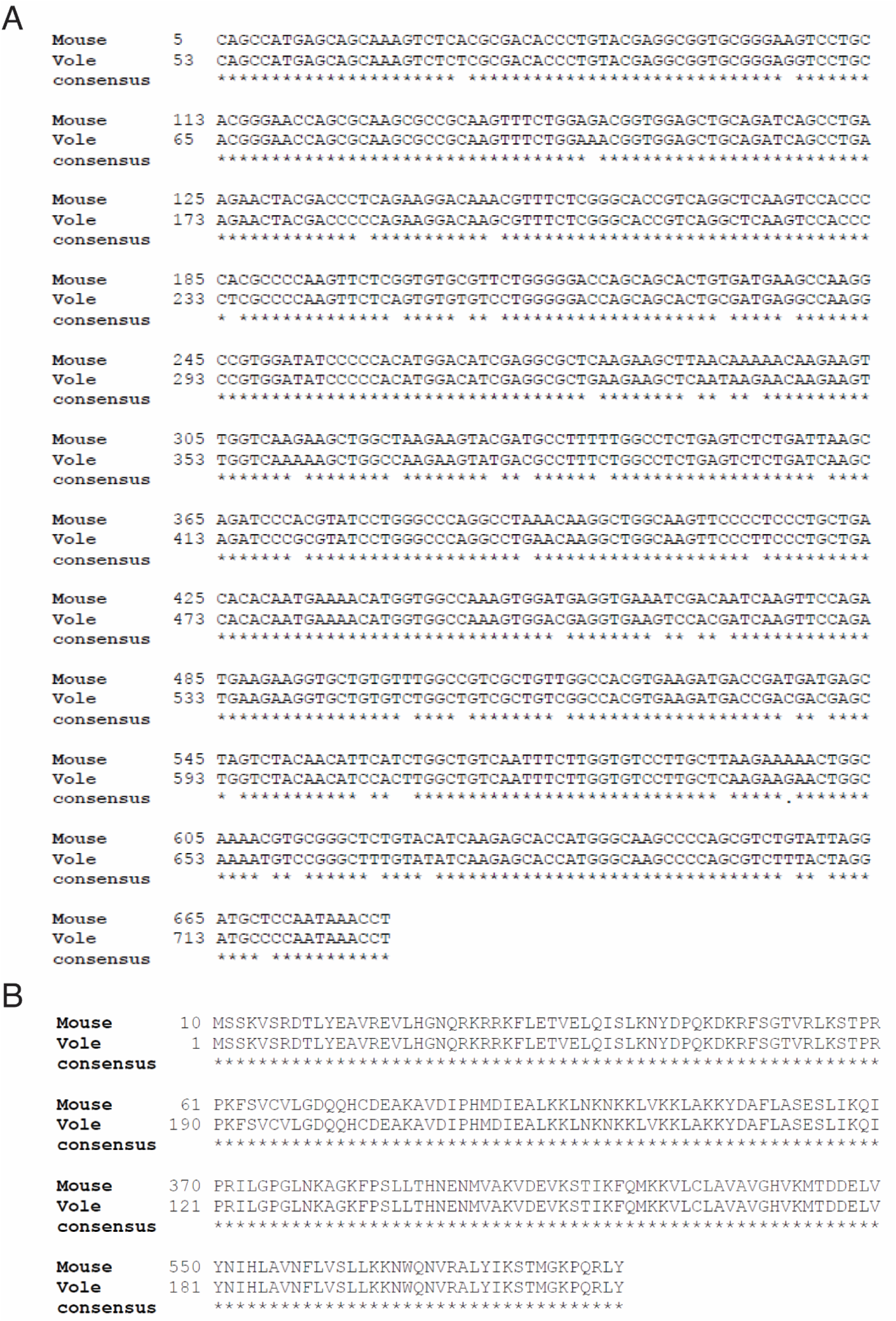
Alignment of mouse (*Mus musculus*) and prairie vole (*Microtus ochrogaster*) ribosomal protein L10a (RPL10a) cDNA sequences and amino acid sequences. **(A)** The mouse RPL10a mRNA sequence (GenBank BC083346.1) was aligned to the prairie vole RPL10a cDNA sequence (XP_005360358.1) using BLASTN 2.8.0+33. Asterisks indicate matching nucleotides. The mouse and vole cDNA sequences are 93% identical, with an expected value of 0.0. **(B)** The mouse RPL10a amino acid sequence was deduced from the mouse RPL10a cDNA sequence (GenBank BC083346.1) and aligned to the prairie vole RPL10a amino acid sequence (XP_005360415.1) using BLASTX 2.8.0+33. Asterisks indicate matching amino acids. The mouse and vole amino acid sequences are 100% identical, with an expected value of 3 × 10^−145^.

**Fig. S9.**
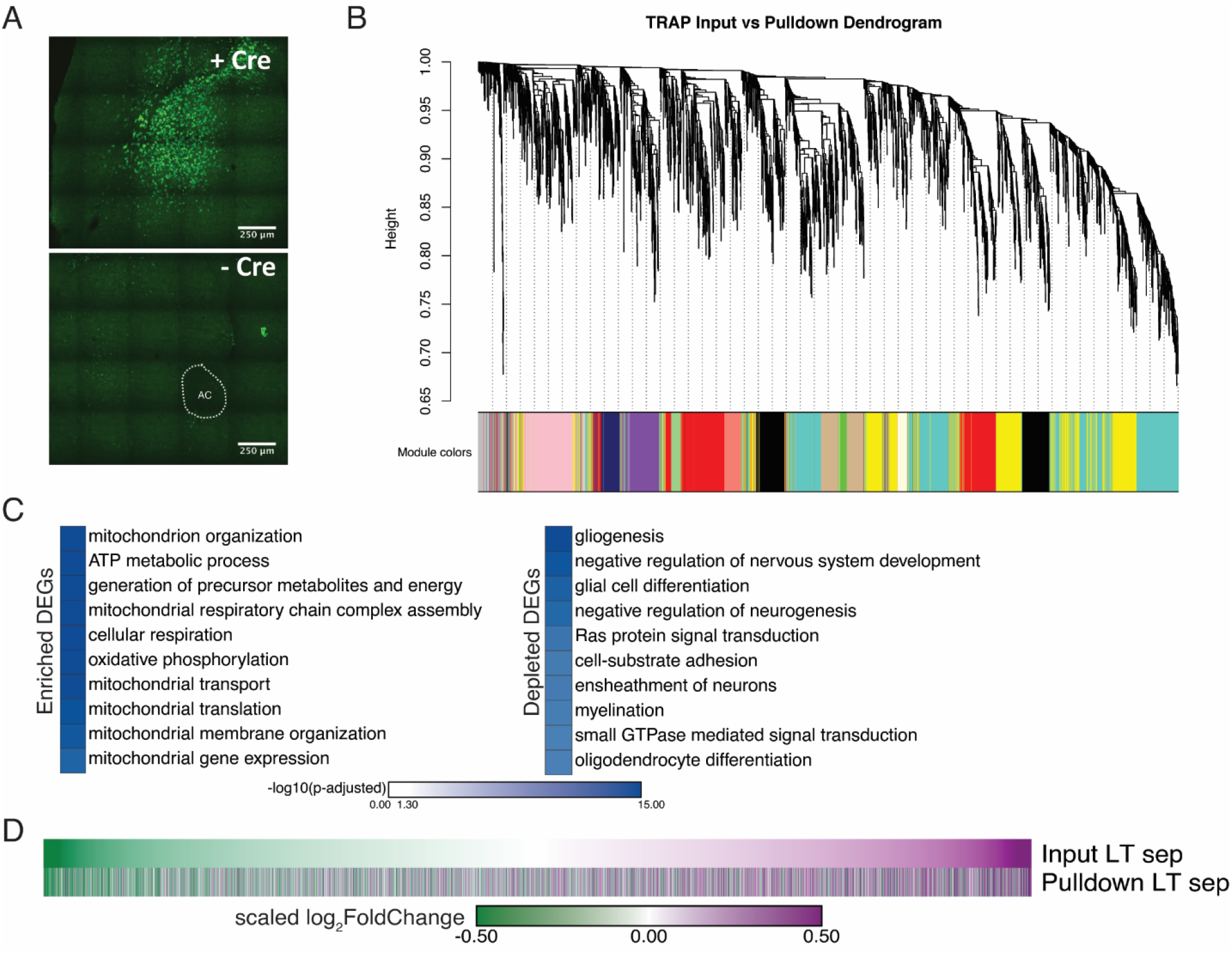
Validation of expression and analyses for vole optimized Translating Ribosome Affinity Purification. **(A)** Vole optimized GFP-RPL10a vector is Cre-dependent as confirmed by bilateral NAc injections +/- Cre. ac = anterior commissure. **(B)** The input and pulldown samples from vTRAP cohorts were used to construct gene co-expression networks using WGCNA. The resulting co-expression networks are clustered in a dendrogram and each gene module is assigned a unique color. **(C)** Gene Ontology analysis using mouse ontology terms indicates that transcripts associated with the mitochondria and cellular respiration are enriched while transcripts associated with gliogenesis and myelination are depleted, supporting neuronal targeting of RPL10-GFP. **(D)** The neuronal enriched genes from *Fig 4E* were used to filter the input and pulldown samples of the opposite-sex 4 week separated cohort. The high correspondence in the scaled log_2_FoldChange of the Morpheus heatmaps validates that, for neuronal enriched genes, their expression in bulk sequencing is largely representative of their expression in neurons.

**Dataset S1 (separate file). Comprehensive statistical analyses and individual animal information**.

Statistical analyses for behavior and transcription experiments including applicable post-hocs and version numbers of analysis packages used. Also includes relevant metrics for all experimental animals including information about excluded animals.

**Dataset S2 (separate file). DESeq, RRHO, and Fisher’s Exact Test intersection gene lists**.

Workbook includes all gene lists used in analysis. For DESeq, all results dataframes and differentially expressed genes are included with all output metrics. RRHO tabs include genes in each list used to generate heatmaps and overlapping genes for each quadrant as well as genes in the Fisher’s Exact Test intersections. Descriptions of all column names, the associated figure, and versions of packages used are in the last three tabs.

**Dataset S3 (separate file). Comprehensive Ingenuity Pathway Analysis data**.

Comprehensive output from IPA analyses for both Canonical Pathways and Upstream Regulators. Includes all terms with those that are statistically significant highlighted in red.

